# Epigenetic and transcriptional regulation of the human angiotensinogen gene by high salt

**DOI:** 10.1101/2023.11.22.568343

**Authors:** Sravan Perla, Ashok Kumar

**Affiliations:** Department of Pharmacology, Yale School of Medicine, New Haven, CT, USA; Department of Pathology, New York Medical College, Valhalla, NY, USA; Department of Physiology and Pharmacology, University of Toledo College of Medicine and Life Sciences, Toledo, OH, USA

**Keywords:** Hypertension, Epigenetics, Transcriptional regulation, Haplotype, RNA-Seq, Liver, Kidney

## Abstract

Hypertension is caused by a combination of genetic and environmental factors. Angiotensinogen (AGT) is a component of RAAS, that regulates blood pressure. The human angiotensinogen (hAGT) gene has -6A/-6G polymorphism and -6A variant is associated with human hypertension. In this study, we have investigated the epigenetic regulation of the hAGT. To understand transcriptional regulation of the hAGT, we have made transgenic animals containing -6A. We show that HS affects DNA methylation and modulates transcriptional regulation of this gene in liver and kidney. High salt (HS) increases hAGT gene expression in -6A TG mice. We have observed that the number of CpG sites in the hAGT promoter is decreased after HS treatment. In the liver, seven CpG sites are methylated whereas after HS treatment, only three CpG sites remain methylated. In the kidney, five CpG sites are methylated, whereas after HS treatment, only three CpG sites remain methylated. These results suggest that HS promotes DNA demethylation and increasing AGT gene expression. RT-PCR and immunoblot analysis show that hAGT gene expression is increased by HS. Chip assay has shown that transcription factors bind strongly after HS treatment. RNA-Seq identified differentially expressed genes, novel target genes associated with hypertension, top canonical pathways, upstream regulators. One of the plausible mechanisms for HS induced up-regulation of the hAGT gene is through IL-6/JAK/STAT3/AGT axis.

## Introduction

Hypertension is a complex disease caused by various genetic and environmental factors (1). Hypertension can lead to renal diseases, metabolic disease, cardiovascular diseases, stroke, heart failure and eventually death (2). Hypertension is defined as the systolic blood pressure over 140 mm Hg or diastolic blood pressure over 90 mm Hg or both (3). Previous studies have shown that excess dietary sodium (Na+) is a major risk factor for hypertension and cardiovascular disease (4–6). The renin–angiotensin-aldosterone system (RAAS) plays a pivotal role in the regulation of blood pressure (7). Angiotensin-II is one of the most active vasopressor agents and is obtained by proteolytic cleavage of angiotensinogen (AGT), which is primarily synthesized by the liver and, to a lesser extent in other tissues such as kidney, adrenal gland, heart, fat, brain and vascular walls (8). The AGT gene is associated with essential hypertension (9–11), cardiac hypertrophy (12), cerebral damage, renal tubular dysgenesis (13, 14) and atherosclerosis (15).

Recent studies have implicated inflammation play an important role in hypertension (6, 16–19). The hAGT gene has -6A/G and -217A/G polymorphism, and -6A/-217A allele is associated with increased blood pressure compared to -6G/-217G (9). We have also observed that the human AGT gene (14 kb) has about 20 SNPs. Variants, -1670A, -1562C, -1561T and 1164A almost always occur with -6A, named it as haplotype-I (Hap -6A) (20). Previous studies have shown that AGT gene expression is influenced by epigenetic modifications such as DNA methylation (21). Epigenetics is defined as the study of the cellular and physiological phenotypic trait variations which are caused by external factors that regulate the gene expression without modifying the underlying DNA sequence (22). DNA methylation occurs at cytosine (C) residues at CpG island context. The CG sites are short regions of DNA where a cytosine is followed by a guanine nucleotide in the linear sequence of bases along its 5’ → 3’ direction. Cytosine base in CpG island can be methylated to form 5-methylcytosine. In mammalian somatic tissues, 70%-80% of CpG islands are methylated (23, 24). During hypomethylation transcription factors and RNA polymerase complex can easily bind to the promoter region and upregulates the expression of the gene (25, 26) On the other hand, hypermethylation is associated with lesser gene expression where DNA methylation blocks the binding of transcription factors and RNA polymerase complex to the promoter of the gene (27).

DNA methylation affects AGT gene regulation and are involved in the alteration of the vascular structure and function in response to environmental stress and diet (28). There is now a growing body of evidence bolstering the role for dietary salt in regulating blood pressure. In this study, we have focused on the epigenetic regulation of the human AGT gene and the effect of high salt on the transcriptional regulation of this gene. Previous studies have shown that excess dietary salt intake is a risk factor for hypertension (29, 30). Our results provide evidence for the first time that high salt induces the DNA demethylation of the hAGT promoter and facilitates the binding of various transcription factors such as USF1, STAT3, HNF1α, C/EBP-β, GR and Pol II to the promoter region and thus upregulate the hAGT gene expression and this may contribute to increased blood pressure.

## Results

### High-salt treatment increases DNA demethylation and upregulates hAGT gene expression in the liver and kidney of -6A TG mice as compared to basal -6A TG mice

To determine the effect of HS on methylation, we treated single transgenic mice containing -6A with 4% high-salt. AGT is primarily synthesized in the liver and is most important contributor of plasma AGT. Therefore, we examined the expression of hAGT in liver of our -6A TG mice in response to HS. As seen in (**Fig.1A**) we found that -6A liver has seven methylation sites. After HS treatment, only three methylation sites were observed in the liver of -6A TG mice (**Fig.1B**). Therefore, liver of HS treated -6A TG mice has higher hAGT gene expression compared to non-treated -6A TG mice. Since AGT is also synthesized in the kidney, we examined the expression of hAGT in kidney of our -6A TG mice in response to HS. As seen in (**Fig.1C**) we found that kidney of -6A TG mice has five methylation sites in basal conditions. After HS treatment, only three methylation sites were observed in the kidney of -6A TG mice (**Fig.1D**). Therefore, HS treated kidney of -6A TG mice has higher hAGT gene expression compared to non-treated kidney of -6A TG mice. High salt induces demethylation, which facilitates the binding of transcriptions factors and RNA polymerase complex to the promoter region and increases gene expression. We found increased number of CpG sites in the liver and kidney of -6A TG mice as compared to high salt treated liver and kidney of -6A TG mice. Therefore, HS treated -6A TG mice have higher AGT gene expression as compared to non-treated -6A TG mice.

**Fig.1.**
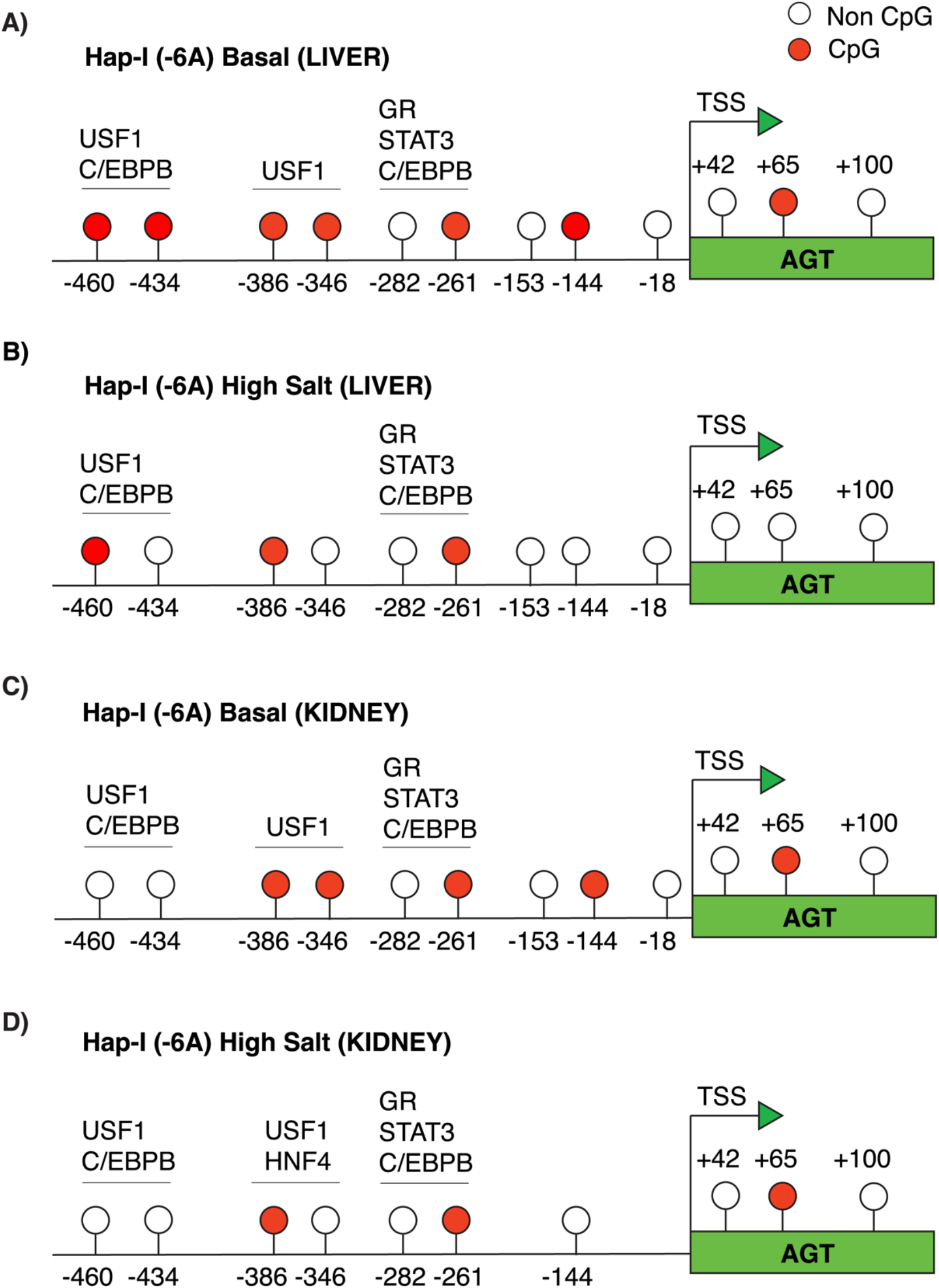
Methylation pattern of the hAGT promoter in the liver and kidney of Hap-I(6A) TG mice after high salt treatment. Seven CpG sites were observed in the liver of Hap-I TG mice (Fig.1A). After high salt treatment four CpG sites were lost and retains three CpG sites (Fig.1B). On the other hand, five CpG sites were observed in the kidney of Hap-I TG mice (Fig.1C). After high salt treatment two CpG sites were lost and retained three CpG sites (Fig.1D). Therefore Hap-I(-6A) liver and kidney has higher hAGT gene expression after high salt treatment compared with Hap-I(-6A) basal liver and kidney.

### High salt increases the hAGT mRNA expression in liver and kidney of -6A TG mice

To determine the effect of methylation on the transcriptional regulation of the hAGT gene we analyzed the hAGT mRNA expression. As seen in (**Fig.2A**) we found that mRNA expression of hAGT is increased by 2.8-fold in the liver of -6A (p<0.05) after high salt treatment. mRNA expression of hAGT is increased by 2-fold in kidney of -6A (p<0.05) after high salt treatment (**Fig.2C**). Whereas endogenous mouse angiotensinogen (mAGT) expression was not significantly different in both liver and kidney tissues (**Fig.2B and D**).

**Fig.2.**
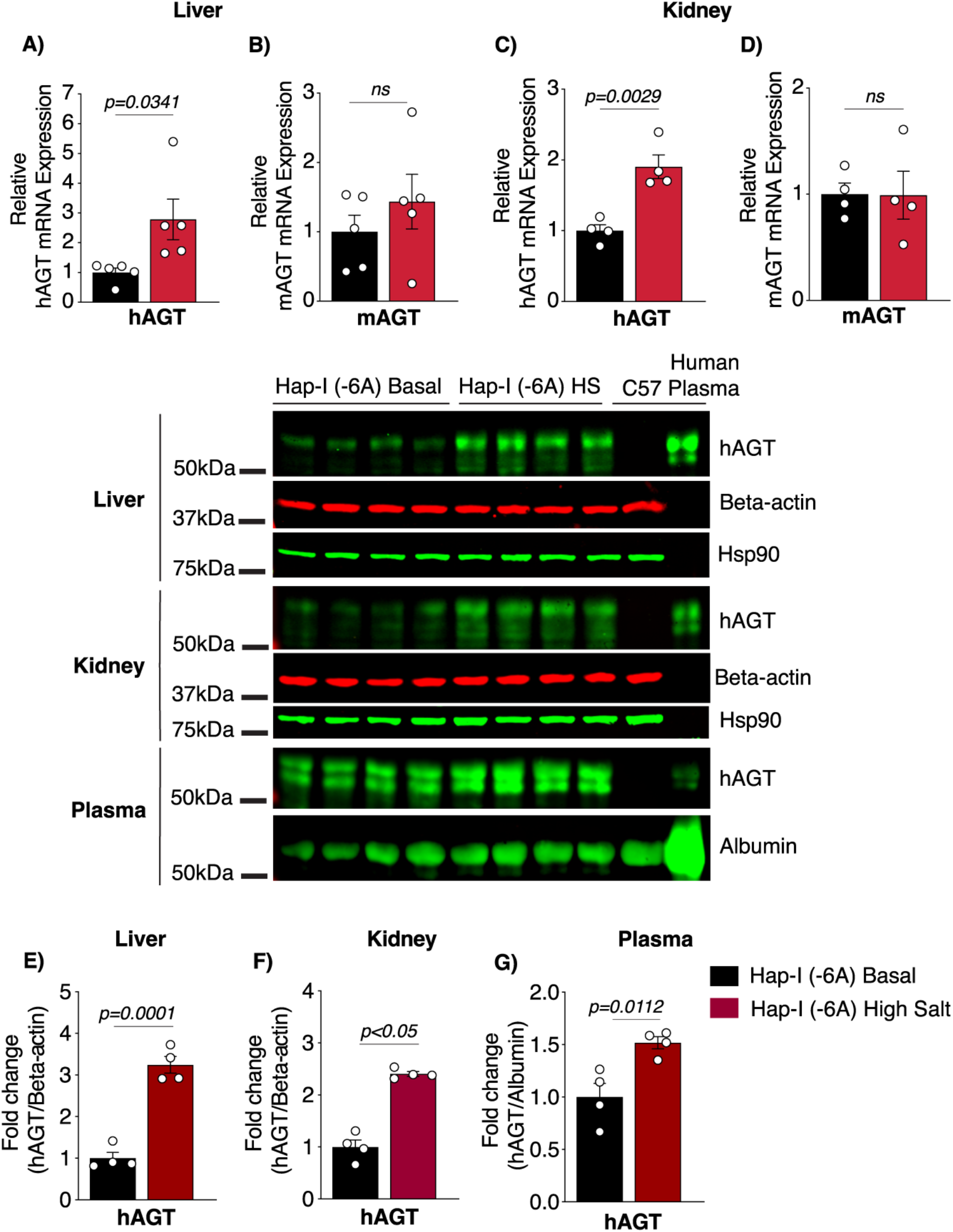
High salt increases hAGT mRNA and protein expression in liver and kidney. The relative mRNA expression levels of hAGT and endogenous mAGT in the liver (n=5) and kidney (n=4) tissues of Hap-I transgenic mice were measured by qRT-PCR. Liver (n=4), kidney (n=4) and plasma (n=4) lysates from 12-week-old male Hap-I(-6A) TG mice were immunoblotted with anti hAGT, beta actin, HSP90 and albumin antibodies. mRNA expression of the hAGT gene (Fig.2A**&C**) in Hap-I TG mice is increased after HSD in both liver and kidney, whereas mRNA expression of endogenous mAGT (Fig.2B**&D**) levels are not significantly different after HSD treatment in both liver and kidney. Immunoblot analysis showing the expression of the hAGT protein is significantly elevated in the liver (Fig.2E), kidney (Fig.2F) and plasma (Fig.2G) of the TG mice with Hap-I (6A) HSD compared with Hap-I basal TG mice. Statistical significance was analyzed with Student’s t-test. All data represent mean ± SEM (*p<0.05).

**Fig.3.**
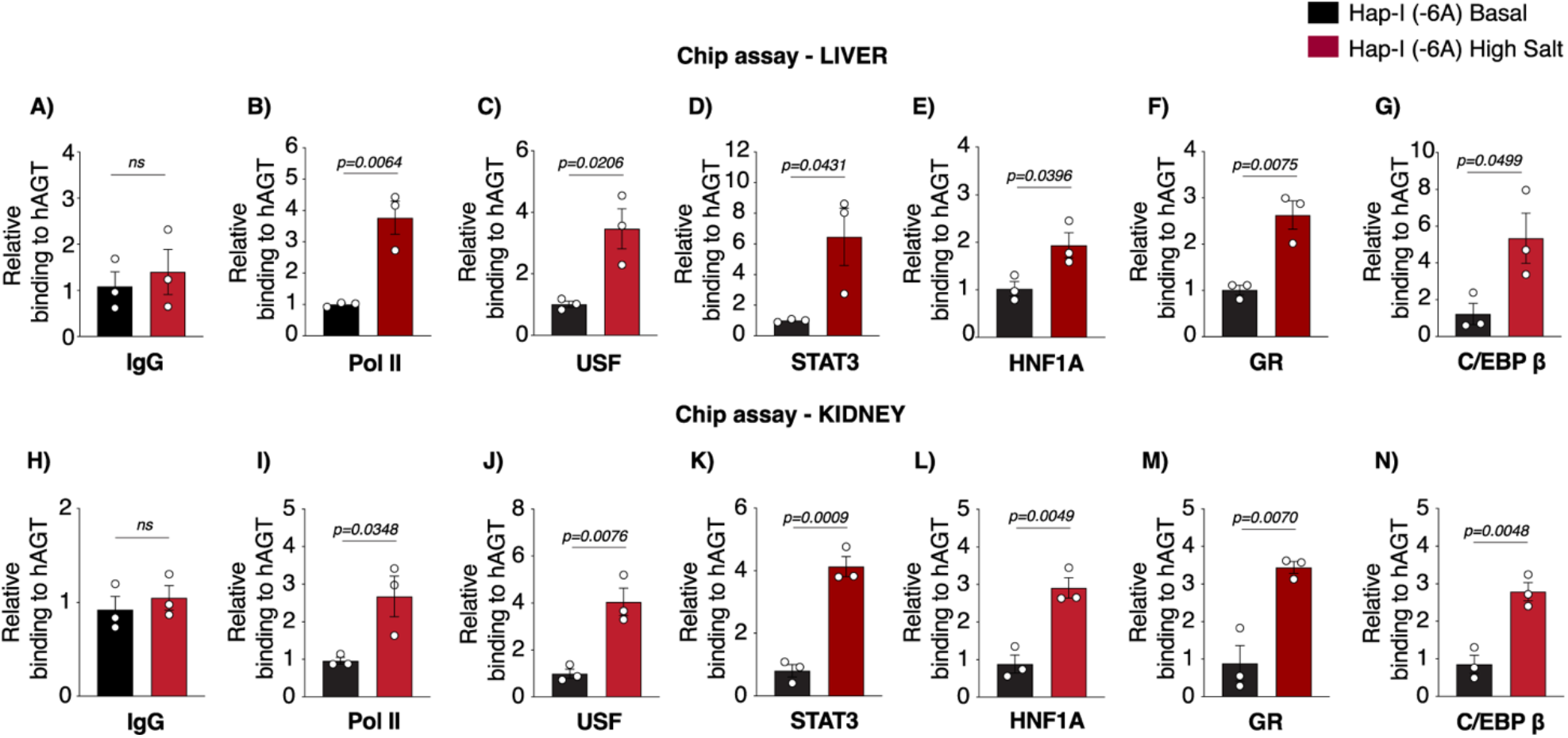
High salt increases the binding of gene regulatory proteins to the hAGT gene promoter. Chip assay showing the relative binding of various transcription factors to the hAGT gene promoter region. The assay was performed in the presence of antibodies against IgG, Pol II, USF, STAT3, HNF1a, GR and C/EBP-β. High salt diet significantly enhances the enrichment of Pol II, USF, STAT3, HNF1a, GR and C/EBP-β in the promoter of the hAGT gene from liver (**Fig.3A-G**) and kidney (**Fig.3 H-N**) of Hap-I(-6A) TG mice. All data represent mean ± SEM. Statistical significance was analyzed with Student’s t-test (*p<0.05).

### High-salt increases hAGT protein levels in the liver, kidney and plasma of -6A TG mice

To determine the effect of methylation on the protein expression of the hAGT gene we analyzed the hAGT protein expression by western blot. As seen in **Fig.2E**, **F** and **G** we found that HS induces hAGT protein expression in the liver (**Fig.2E**) kidney (**Fig.2F**) and plasma (**Fig.2G**) of transgenic mice with -6A (*p<0.05). Protein expression of hAGT is increased by 3-fold in the liver (**Fig.2E**) and 2.3-fold in the kidney (**Fig.2F**) after high salt treatment. hAGT protein expression is increased by 1.6-fold in the plasma after high salt treatment (**Fig.2G**).

### Chip assay shows that high salt treatment increases binding of USF1, STAT3, HNF1a, GR and C/EBP-β to the promoter of hAGT gene and upregulates it’s expression

We next examined the effect of HS on the binding of various transcription factors at different locations on the hAGT promoter. Previous studies also have shown several transcription factors such as STAT3, HNF1a, USF, GR and C/EBP-β binds to the hAGT promoter (31). However, we wanted to investigate whether altered binding of any transcription factors involved in this process. We examined the effect of HS on the binding of STAT3 to the promoter region of the hAGT gene in - 6A TG mice. As shown in **Fig.4D** and **K** we found that HS treatment significantly enhances the enrichment of STAT3 in -6A TG mice. These data suggest that HS treatment promotes the binding of STAT3 to the promoter of hAGT gene. This leads to up-regulation of hAGT gene in the liver and kidney of these TG mice. We examined the effect of HS on the binding of USF-1 to the chromatin region of the hAGT gene in -6A transgenic mice. As shown in **Fig.4C** and **J**, we found that HS treatment significantly enhances the enrichment of USF-1 to the chromatin around the region of -460 in the transgenic mice with -6A. These data suggest that HS promotes the binding of USF-1 to the promoter of the hAGT gene. This leads to up regulation of hAGT gene in the liver and kidney of -6A TG mice. Transcription factor C/EBP-β has been shown to bind to the hAGT promoter region and regulate it’s expression by inducing chromatin relaxation (32). We found HS increases the binding of C/EBP-β to the hAGT promoter region in -6A TG mice and upregulates its expression (**Fig.4G** and **N**). We have earlier shown that HNF-1α increase the expression of the hAGT gene (33). Chip assay has shown that HS increases the binding of HNF-1 alpha to the hAGT promoter region in -6A TG mice. One of the possible mechanisms could be HS induces the chromatin relaxation by DNA demethylation. This leads to the binding of HNF-1alpha to the hAGT promoter and upregulates its expression (**Fig.4E** and **L**). HS also increases the binding of GR to the hAGT promoter region and upregulates hAGT gene expression in -6A TG mice (**Fig.4F** and **M**). We have noticed that high salt increases the binding of Pol II to the hAGT promoter in this haplotype (**Fig.4B** and **I**). Pol II and IgG acts as a positive and negative control respectively. We have observed that STAT3 binds to hAGT promoter region more strongly than other transcription factors. In conclusion our chip assay shows that HS increases the binding of transcription factors USF1, STAT3, HNF-1α, GR and C/EBP-β to the promoter of hAGT region and upregulates it’s expression.

**Figure.4.**
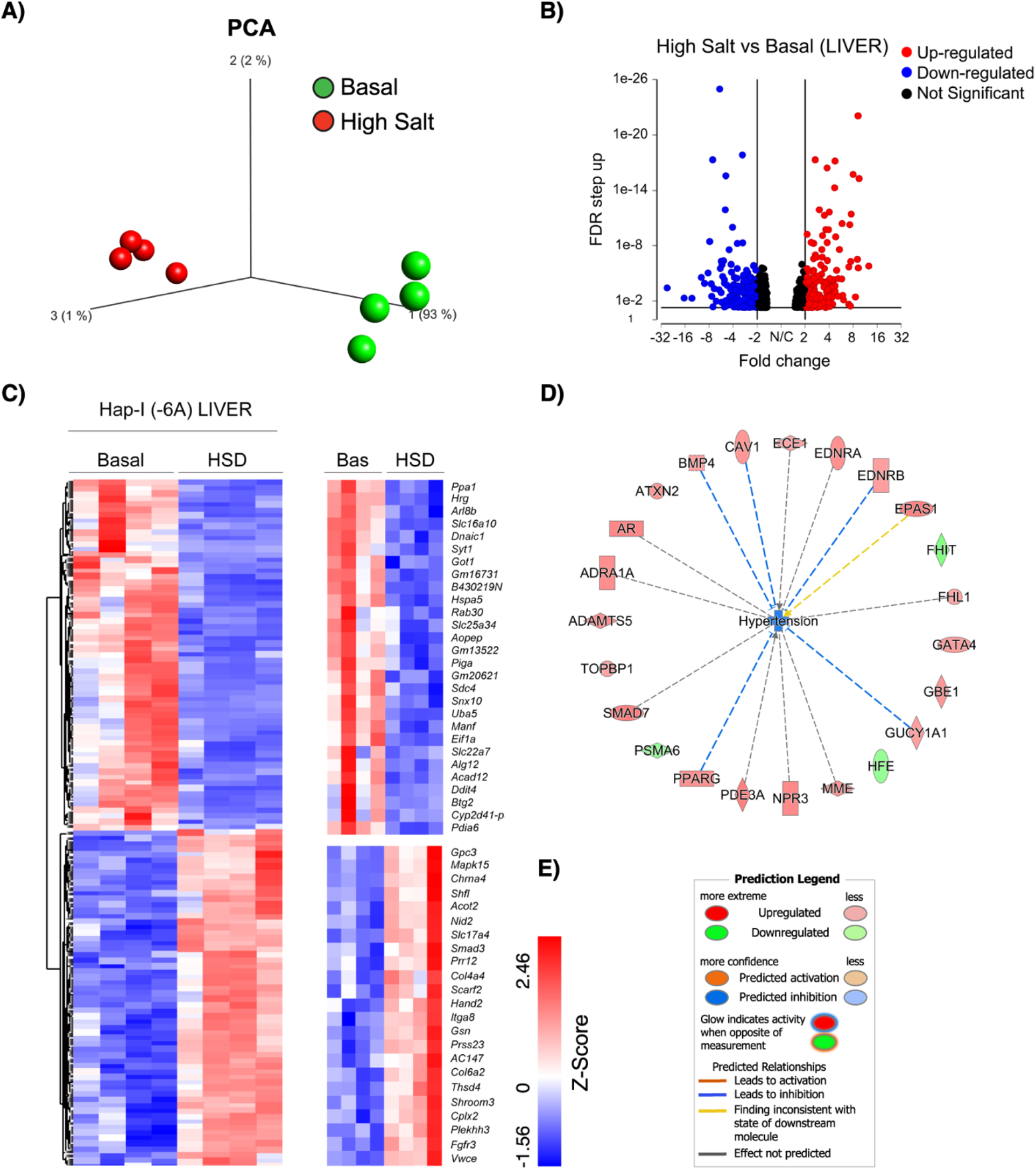
RNA-Seq whole transcriptome analysis in the liver of high salt treated Hap-I TG mice. Total RNA was isolated from the liver of 12-week-old male TG mice and RNA seq analysis was performed. Principal component analysis plot (PCA) displaying two groups along PC1, PC2 and PC3 which describe 1%, 2% and 93% of the variability respectively (**Fig.4A**). PCA analysis was applied to normalized counts and log-transformed count data. Volcano plot of differentially expressed genes in basal and high salt treated Hap-I TG mice (**Fig.4B**). The expression difference is considered significant for a fold change ≤ -1.5 or ≥1.5 (FDR step up ≤ 0.05). Downregulated genes are shown in blue and upregulated genes are shown red in color. Hierarchical clustered heatmap of log2-transformed gene expression (FDR step up ≤ 0.05) in the liver of Hap-I TG mice. Each row represents a gene, and each column represents an individual mouse. Differential gene expression was shown in the heatmap (**Fig.4C**). Ingenuity pathway analysis (IPA) performed on statistically significant genes to identify high salt responsive genes involved in hypertension (**Fig.4D**).

### Transcriptome analysis in the liver and kidney of -6A transgenic mice after HS treatment

To provide insight into the effects of HS on the overall gene expression profile in the -6A TG mice, whole-transcriptome RNA-sequencing (RNA Seq) was performed in liver and kidney tissues of these mice. We have generated approximately 22-49 million reads per sample, filtered reads to have high quality scores. We mapped 84.1%–86.3% of those reads to the mouse genome. Principal component analysis (PCA) showed that samples from the basal and high salt clustered separately, suggesting that transcriptome profiles were different after HS treatment (**Fig.4A** and **6A**). We have identified differentially expressed genes (liver; FDR ≤ 0.05, fold change ≥1.5 and kidney; *p*<0.05, fold change ≥1.5): 987 genes were found to be differentially expressed in the liver between normal and HS, with 502 downregulated and 485 upregulated as shown in the volcano plot (**Fig.4B**). A total of 372 genes were found to be differentially expressed in the kidney between basal and HS, with 175 downregulated and 197 upregulated as shown in the volcano plot (**Fig.6B**).

Ingenuity pathway analysis (IPA) in the **liver** tissue has identified hypertension–related genes such as *Ece1, Ednra, Ednrb, Epas1, Fhit, Fhl1, Gata4, Gbe1, Gucy1a1, Hfe, Mme, Npr3, Pde3a, Pparg, Psma6, Smad7, Topbp1, Adamts5, Adra1a, Ar, Atxn2, Bmp4 and Cav1*. *Fhit, Hfe* and *Psma6* genes were found to be downregulated whereas all the remaining genes were found to be upregulated (**Fig.4D**). *Npr3* encodes natriuretic peptide receptor 3. *Npr3* regulate blood volume, cardiac function, pulmonary hypertension, and abdominal fat distribution (34). *Bmp4* encodes bone morphogenetic protein 4 and belongs to Transforming Growth Factor (TGF)-superfamily. *Bmp4* is upregulated in hypoxia-induced pulmonary hypertension (35). Caveolin-1 (*Cav1*) is a structural component of caveolae in the membrane. *Cav1* is expressed in both vascular smooth muscle cells and endothelium. *Cav1* is implicated in pulmonary hypertension, several cardiovascular diseases including atherosclerosis and dilated cardiomyopathy (36). The hierarchical clustering of log ratio–transformed gene expression was represented in a heatmap and showed genes that were differentially expressed between basal and high salt treated liver of -6A TG mice (**Fig.4C**). Further, these results show that some of these genes (*Ppa1, Hrg, Syt1, Rab30, Sdc4, Ddit4, Btg2, Snx10, Uba5, Manf, Pdia6, Gpc3, Mapk15, Acot2, Smad3, Hand2, AC147, Ita8, Gsn, Shroom3, Thsd4, Fgfr3, Col6a2 and Vwce*) expression reversed after HS treatment (**Fig.4C**). Genes exhibiting revertant behavior (**Fig.4C**) demonstrate that they are haplotype specific and hence represent a transcriptomic “fingerprint” of -6A target genes that are potentially involved in the progression of hypertension. Further we have performed Gene Set Enrichment Analysis (GSEA) (**Fig.5A**) and Pathway Enrichment Analysis (PEA) (**Fig.5B**) in the liver of -6A TG mice. GSEA deals with the groups of genes that share common biological function. In our GSEA analysis, we have identified that many of our genes are linked to the regulation of blood pressure and interestingly it’s showing the highest enrichment score of 4 (**Fig.5A**). PEA helps scientists gain mechanistic insight into gene lists generated from RNA-Seq analysis. PEA also identifies biological pathways that are enriched in a gene list more than would be expected by chance (37). Interestingly, we have found that many of our genes are linked to renin-angiotensin system and JAK-STAT signaling pathway (**Fig.5B**). We also identified top canonical pathways (**Fig.5C**), upstream regulators (**Fig.5D**), top diseases and functions (**Fig.5E**), and top tox functions (**Fig.5F**) in -6A TG mice after HS treatment. “Upstream regulator” refers to any molecule that can affect the expression, transcription, or phosphorylation of another molecule. Interestingly, we have identified *Il-6, Stat3, Irs1, Rgs2*, *Tgf-β* and *Fasn* as some of our top upstream regulators (**Fig.5D**). Tox functions refers to the genes associated with toxicity endpoints, phenotypes, and their causal associations, when known. We have identified pulmonary hypertension as one of our top tox function (**Fig.5F**). In **Fig.5G**, we have mentioned the graphical summary of the different molecules, functions and pathways involved in the pathogenesis of hypertension in the liver of -6A TG mice.

**Figure.5.**
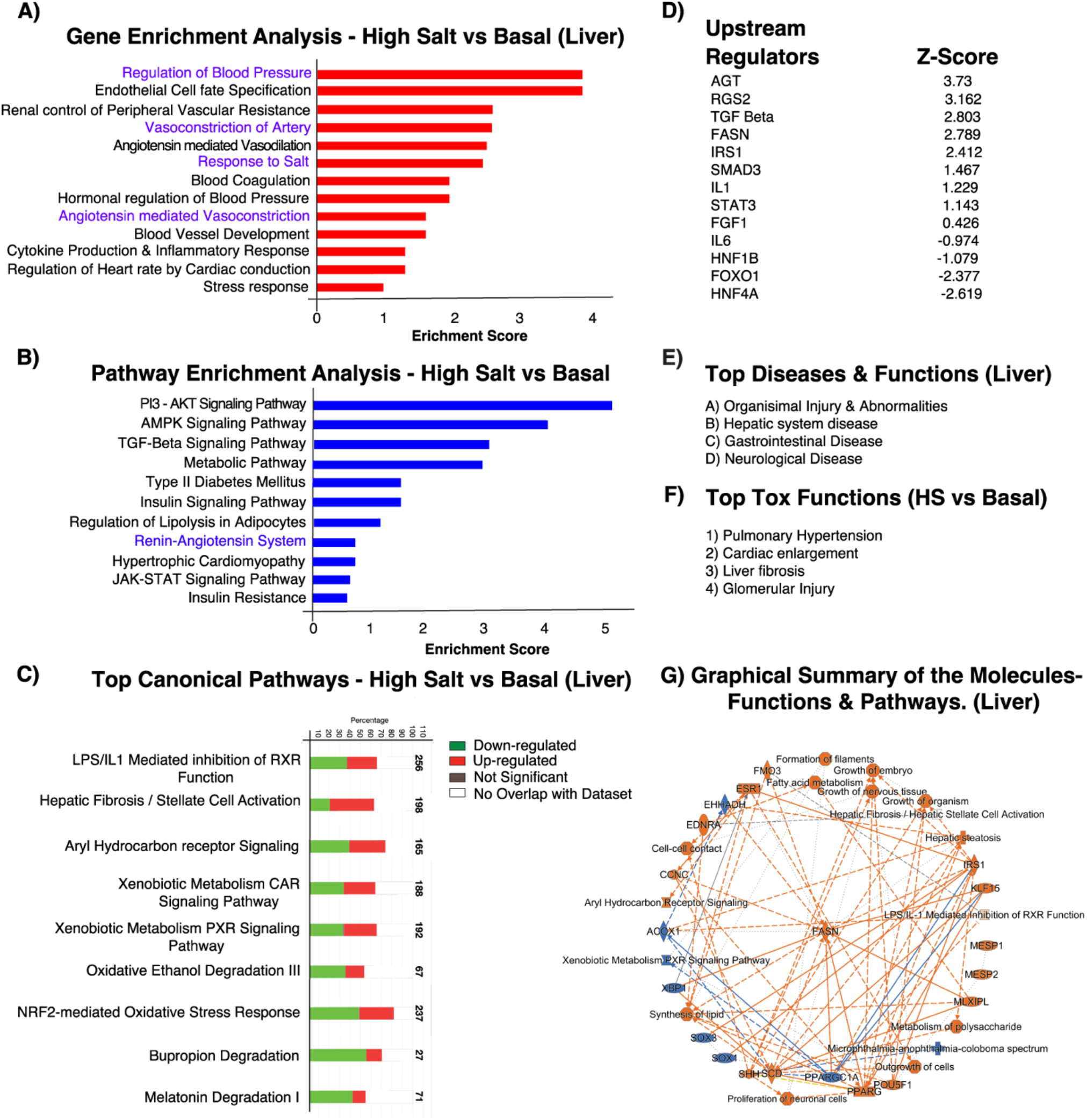
Gene ontology analysis in the liver of high salt treated Hap-I(-6A) TG mice. Gene ontology (GO) enrichment analysis (**Fig.5A**), pathway enrichment analysis (**Fig.5B**) was performed from the gene list of basal and high salt treated Hap-I(-6A) TG mice. Top biological processes were ranked by enrichment score. upstream transcription regulators (**Fig.5D**), top canonical pathways (**Fig.5C**), top diseases & functions (**Fig.5E**), and top tox functions (**Fig.5F**) were analyzed. Graphical summary of the molecules, functions and pathways are mentioned (**Fig.5G**).

IPA also identified alterations in the expression of hypertension-related genes in the **kidney**. These genes include *Ciart, Cndp1, Cyp3a5, Cyp4a11, Dlg2, Fabp1, Fgb, Gabrb3, Gstp1, Il12B, Il1b, Kcnj2, Kcnma1, Mt-rnr2, Myh6, Nqo1, Ptger3, Ren, Serpina3, Tnnt2, Ttn, Alb, Angpt2, Aplnr, Arg2, Atp4a, Ccng1 and Chn2. Ciart, Cndp1, Cyp4a11, Gstp1, Kcnj2, Nqo1, Angpt2, Aplnr and Arg2* genes were found to be up-regulated whereas all the remaining genes were found to be down-regulated (**Fig.6D**). *Cyp3a5* has been implicated in the regulation of blood pressure and may serve as a potential risk factor for the development of hypertension. Increased levels of *Cyp3a5* could cause sodium and water retention by affecting the metabolism of cortisol (38). Interleukin (IL) *Il12B* is an important proinflammatory cytokine and may be involved in a variety of inflammatory diseases, including hypertension (39). NAD(P)H Quinone Dehydrogenase 1 (*Nqo1*) encodes a cytoplasmic 2-electron reductase. Angiotensin-converting enzyme (*ACE*) plays a key role in blood pressure homeostasis. *ACE* is regulated by *Nqo1* (40). The hierarchical clustering of log ratio– transformed gene expression was represented in a heatmap and showed genes that were differentially expressed between normal and high salt treated kidney of -6A TG mice (**Fig.6C**). Further, we have shown that the genes in the heatmap that were apparently reverted in their expression to high salt treatment (**Fig.6C**). Genes exhibiting revertant behavior (**Fig.6C**) demonstrate that they are haplotype specific and hence represent a transcriptomic “fingerprint” of Hap-I or -6A target genes that are potentially involved in the progression of hypertension. Some of the genes show revertant behavior include *Xlr4b, Pcdhb11, Lax1, Gjb1, Id1, Aqp4, Ly6d, Cyp4a14, and Ighm.* Further we have also performed Gene Set Enrichment Analysis (**Fig.7A**) and Pathway Enrichment Analysis (**Fig.7B**) in the kidney of -6A TG mice. In our GSEA analysis, we have identified that many of our genes are linked to the cytokine mediated signaling pathway, renal water reabsorption and regulation of blood pressure by RAAS with an enrichment score of 7.8, 3.8 and 2 respectively (**Fig.7A**). PEA identifies many of our genes are linked to renin secretion and JAK-STAT signaling pathway with an enrichment score of 5.8 and 1.2 respectively (**Fig.7B**). We also identified top canonical pathways (**Fig.7C**), upstream regulators (**Fig.7D**), top diseases and functions, molecular and cellular functions (**Fig.7E**) in -6A TG mice after HS treatment. Interestingly, we have identified *Il-6, Ahr, Pias4, Clec10a, Stat5b, Cebpa* and *Trim21* as some of top upstream regulators (**Fig.7D**). Furthermore, we have identified endocrine system disorders is one of the top diseases (**Fig.7E**). In **Fig.7F**, we have mentioned the graphical summary of the different molecules, functions involved in the pathogenesis of hypertension in the kidney of -6A TG mice. Furthermore, we have identified several molecules from the RNA Seq analysis based on their implication in hypertension. These genes *include Ar, Fgf21, Foxoa1, Hnf1a, Mapk15, Rgs16 and Fasn.* Androgen receptor (*Ar*) is a type of nuclear receptor that functions as a transcription factor. *Ar* has been shown to be associated with hypertension, stroke, atherosclerosis, abdominal aortic aneurysm, and heart failure (41). Fibroblast growth factor 21 (*FGF21*) is a liver-secreted peptide hormone known to be associated with blood pressure (42). *Foxoa1* belongs to the forkhead family of transcription factors which are characterized by a distinct forkhead domain. The specific function of *Foxoa1* has not yet been determined. However, *Foxoa1* may play a role in cell proliferation and transformation (43). Hepatocyte nuclear factor 1 homeobox A (*Hnf1a*) acts as a transcription factor that is highly expressed in the liver and is involved in the regulation of several liver-specific genes. Mutations in the *Hnf1a* gene have been known to cause diabetes and coronary heart disease (44–46). Mitogen-activated protein Kinase (*Mapk15*) involved in cell signaling cascades that regulate proliferation, differentiation, and transcriptional regulation. Regulator of G-protein signaling 16 (*Rgs16*) inhibits signal transduction by increasing the GTPase activity (47). Fatty acid synthase (*FASN*) is a multi-enzyme protein that catalyzes fatty acid synthesis. Fasn is regulated by Upstream Stimulatory Factors (*USF1* and *USF2*). Fasn is associated with hypertension and metabolic dysfunction (48–50). We have validated these molecules by qRT-PCR both in liver and kidney tissues of -6A TG mice. We have observed that *Ar, Fgf21, Foxoa1, Hnf1a, Mapk15, Rgs16* and *Fasn* mRNA expression is elevated after HS treatment in both liver and kidney of -6A TG mice (**Fig.8 A-N**).

**Figure.6.**
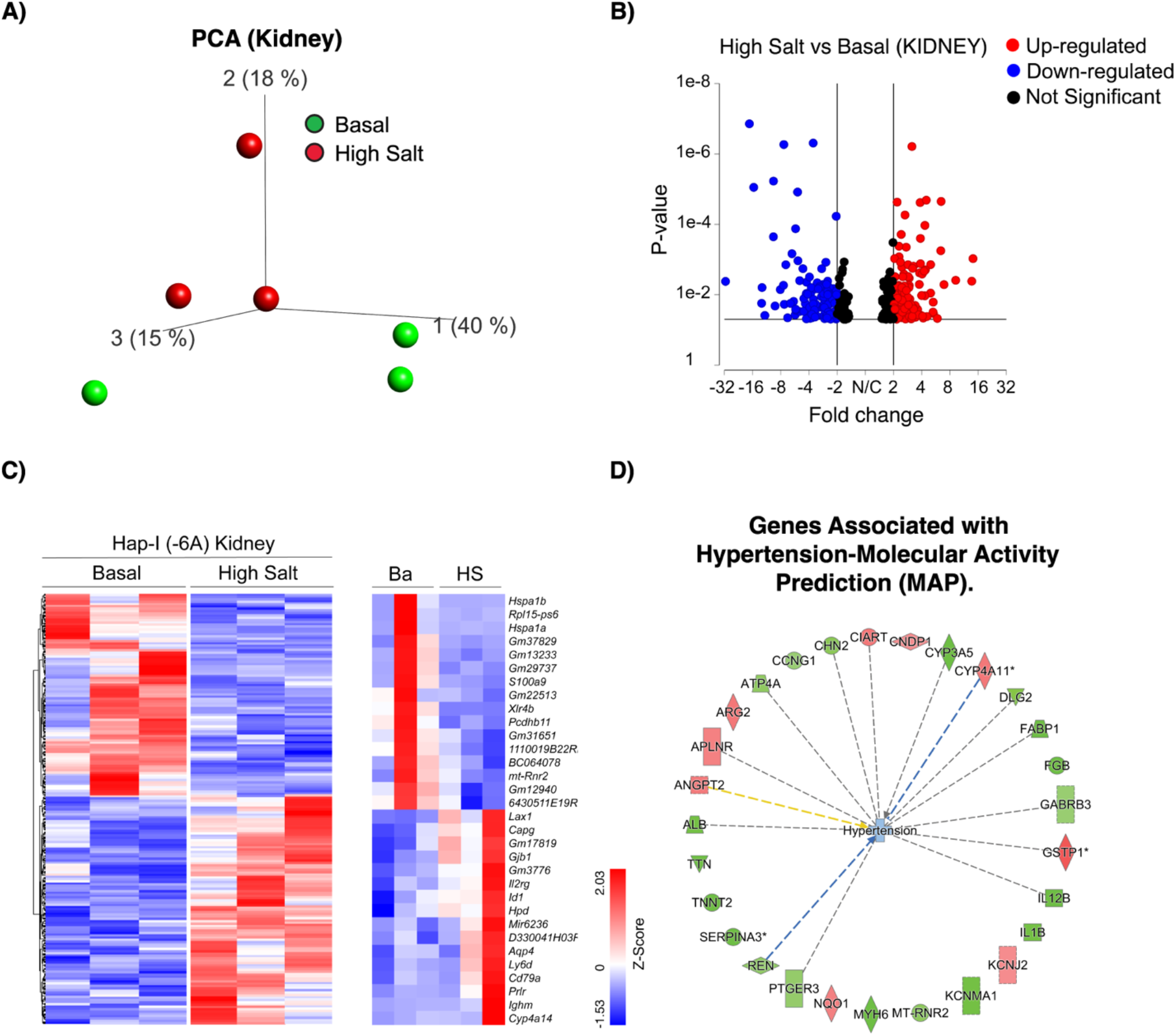
RNA-Seq whole transcriptome analysis in the kidney of high salt treated Hap-I TG mice. Total RNA was isolated from the kidney of 12-week-old male TG mice and RNA seq analysis performed. Principal component analysis plot (PCA) displaying two groups along PC1, PC2 and PC3 which describe 15%, 18% and 40% of the variability respectively. PCA analysis was applied to normalized counts and log-transformed count data (**Fig.6A**). Volcano plot of differentially expressed genes in basal and high salt treated Hap-I TG mice. The expression difference is considered significant for a fold change ≤ -1.5 or ≥ 1.5 (p ≤ 0.05, FDR step up ≤ 0.1). Downregulated genes are shown in blue and upregulated genes are shown red (**Fig.6B**). Hierarchical clustered heatmap of log2-transformed gene expression (p≤ 0.05) in the kidney of Hap-I TG mice. Each row represents a gene, and each column represents an individual mouse. Differential gene expression was shown in the heatmap (**Fig.6C**). Ingenuity pathway analysis (IPA) performed on statistically significant genes to identify high salt sensitive genes involved in hypertension (**Fig.6D**).

**Figure.7.**
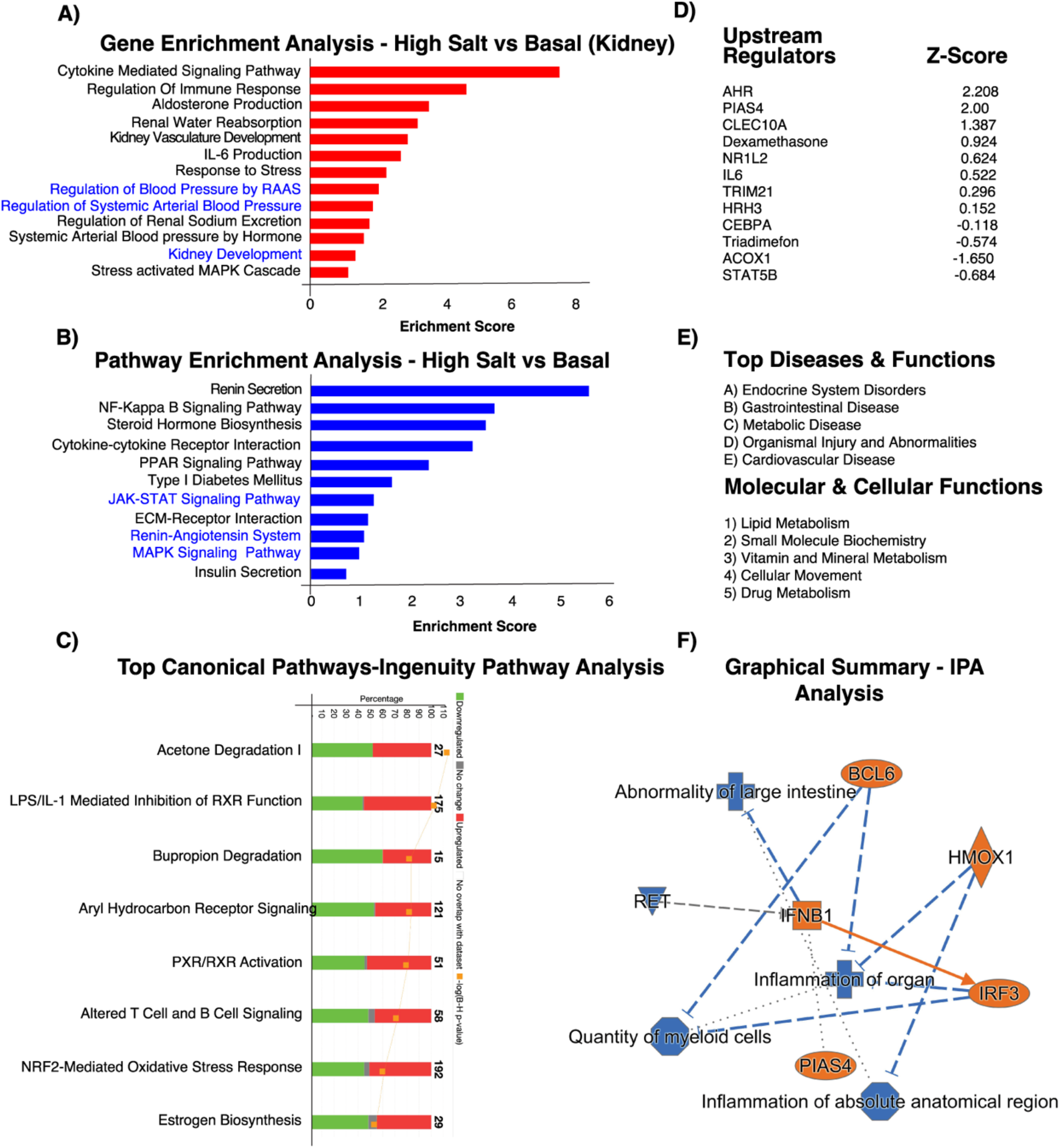
Gene ontology analysis in the kidney of high salt treated Hap-I TG mice. Gene ontology (GO) enrichment analysis (**Fig.7A**) and pathway enrichment analysis (**Fig.7B**) was performed from the gene list of basal and high salt treated Hap-I TG mice. Top biological processes were ranked by enrichment score. Upstream transcription regulators (**Fig.7D**), top canonical pathways (**Fig.7C)** top diseases & functions and molecular and cellular functions (**Fig.7E**) were analyzed. Graphical summary of the molecules and functions are analyzed (**Fig.7F**).

**Figure.8.**
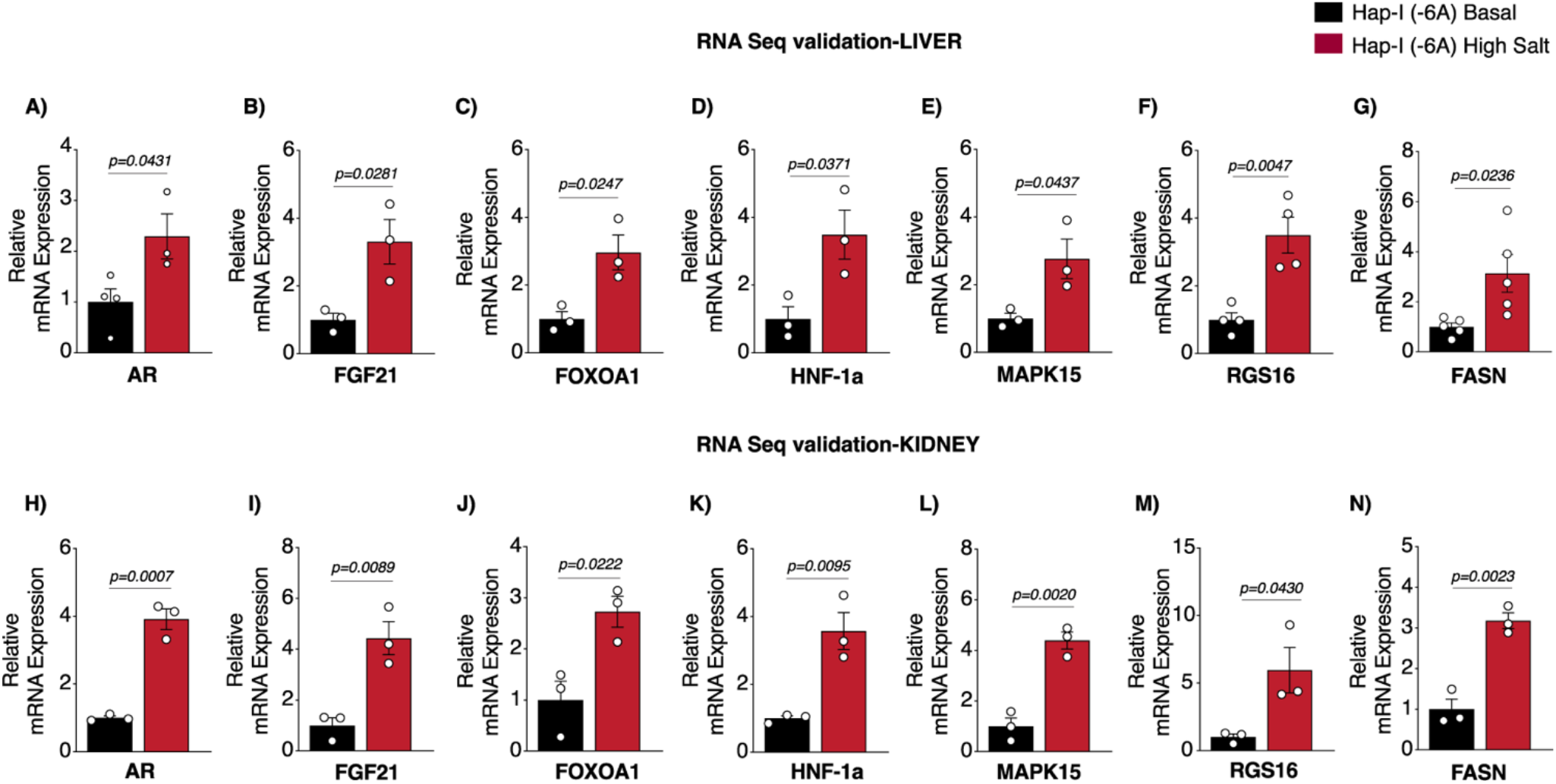
Validation of RNA Seq derived genes by qPCR. Total RNA was isolated from the liver and kidney of 12-week-old male mice and cDNA was prepared and performed qRT-PCR. The relative mRNA expression levels of Ar, FGF21, FOXOA1, HNF1a, MAPK15, RGS16 and FASN in the liver and kidney tissues of Hap-I transgenic mice were measured by qRT-PCR. mRNA expression of the AR, FGF21, FOXOA1, HNF1a, MAPK15, RGS16 and FASN genes in Hap-I TG mice are increased after HSD in both liver (**Fig.8A-G**) and kidney (**Fig.8H-N**). Statistical significance was analyzed with Student’s t-test. All data represent mean ± SEM.

**Figure.9.**
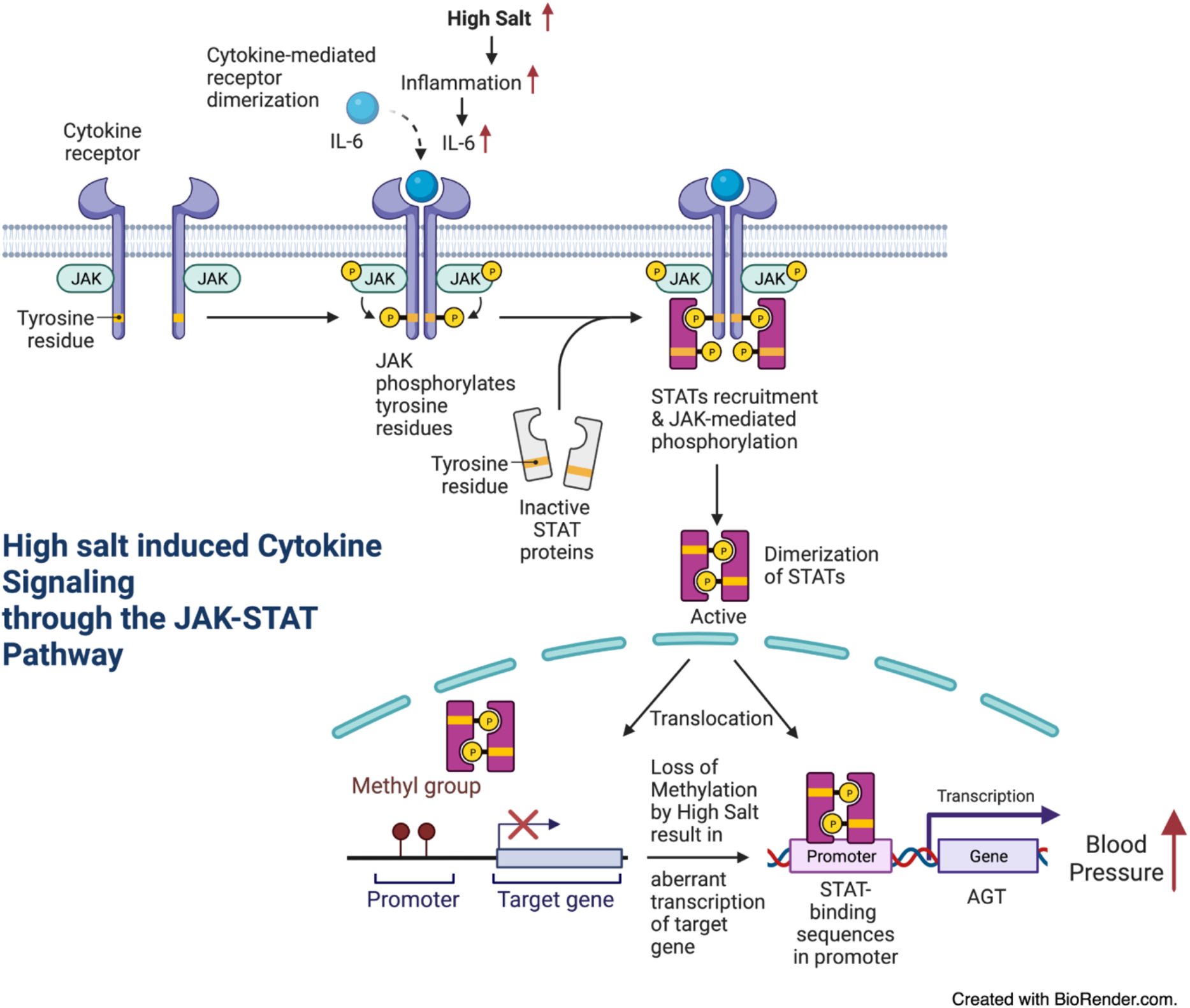
Hypothetical cartoon of high-salt diet induced hAGT gene expression. High salt diet induces inflammation which increases the production of inflammatory cytokines such as IL-6. Upon binding to the cytokine receptor, JAKs add phosphate groups to the cytokine receptor. STAT proteins bind to these phosphate groups on the cytokine receptor and JAK further phosphorylates STATs and forms STAT dimer. Dimerized STAT enters nucleus and binds to the promoter of AGT gene and may increases gene expression (**Fig.9**).

Interestingly, transcriptomic analysis by RNA-seq in the liver and kidney of our -6A TG mice has identified several novel target genes (*Ar, Fgf21, Foxoa1, Hnf1a, Mapk15, Rgs16* and *Fasn*) may implicate in the pathogenesis of hypertension. Transcriptomic analysis contributes to a change in the holistic understanding of hypertension. It enables an insight into the molecular mechanisms underlying hypertension, which has given rise to numerous trials related to transcriptome-based precision medicine.

## Discussion

In this study, we have identified a novel mechanism by which HS treatment contributes to inflammation and hypertension. Our results indicate that HS treatment promotes DNA demethylation, there by increases the binding of transcription factors to the hAGT promoter and modulates hAGT gene expression. Increased AGT gene expression may contribute to inflammation and hypertension. HS is one of the important risk factors in the development and maintenance of hypertension (6, 30). GWAS (Genome Wide Association Studies) studies have linked several alleles with high blood pressure. However, the mechanisms involved in the pathophysiology of hypertension remains elusive (51).

In our previous studies, we have shown that Hap-I (-6A and -217A) of the hAGT gene is associated with hypertension in Caucasians. To understand the role of these genetic variants of the hAGT gene in hypertension, and other cardiovascular abnormalities, we have generated TG mice containing -6A of the hAGT gene. Using these novel -6A TG mice model, we show here, for the first time, that high salt alters DNA methylation pattern, further the transcriptional milieu of cells with up-regulation of the hAGT gene.

The important observation in this study is that HS promotes DNA demethylation thereby positively regulates the hAGT gene expression. High salt upregulated hAGT expression in both liver and kidney of -6A TG mice. Previous studies have shown that diet induces the hAGT gene expression in liver (52–55). However, this is the first study showing that SNP differences in the haplotype-I(-6A) upregulate it’s expression in response to high-salt treatment. Several transcription factors have been implicated in regulation of hAGT gene expression (56, 57). Of these, there is an accumulating body of data supporting the role of several transcription factors such as C/EBP-β, GR, STAT3, HNF-1α and USF1. Effect of HS on DNA methylation and on the transcription factors such as STAT3, C/EBP-β, USF1, GR and HNF-1α was not completely understood. Interestingly, our results show that HS promotes the binding of these transcription factors such as HNF-1α, STAT3, C/EBP-β, GR and USF1 to the hAGT gene promoter and upregulates it’s expression.

HS is one of the key factors in the development and maintenance of hypertension (58). High sodium intake impairs endothelial function and may contribute blood pressure, although the exact mechanisms remain elusive (30). HS induces the susceptibility to adipogenesis and hypertension by altering the methylation pattern, and through the MAPK-STAT3 pathway (59, 60). Inflammation induces the levels of interleukin 6(IL-6) which further binds to IL-6 receptor and activate the Janus kinase/signal transducer and activator of transcription 3 (JAK/STAT3) signaling pathway (61). JAK phosphorylate and activate STAT3, which in turn translocate into nucleus and bind to AGT gene and up-regulate the AGT gene expression. In the present study we observed that SNPs in the promoter of HS treated -6A of the hAGT gene have stronger binding with HNF-1α compared with -6A basal. In conclusion, high salt treatment promotes DNA demethylation which leads to increased chromatin accessibility to various transcription factors. Binding of various transcription factors to the demethylated AGT promoter region increases AGT gene expression. Further, high salt treatment increases inflammation, which leads to the elevation of inflammatory cytokines such as IL-6. Binding of IL-6 to the IL-6 receptor, activates JAK-STAT pathway and finally STAT3 binds to the AGT promoter region and upregulate its expression.

Transcriptome analysis identifies several novel target genes which may implicated in the pathogenesis of hypertension. These genes include *Ece1, Ednra, Ednrb, Epas1, Fhit, Fhl1, Gata4, Gbe1, Gucy1a1, Hfe, Mme, Npr3, Pde3a, Pparg, Psma6, Smad7, Topbp1, Adamts5, Adra1a, Ar, Atxn2, Bmp4, Cav1, Ciart, Cndp1, Cyp3a5, Cyp4a11, Dlg2, Fabp1, Fgb, Gabrb3, Gstp1, Il12B, Il1b, Kcnj2, Kcnma1, Mt-rnr2, Myh6, Nqo1, Ptger3, Ren, Serpina3, Tnnt2, Ttn, Alb, Angpt2, Aplnr, Arg2, Atp4a, Ccng1, Chn2, Ar, Fgf21, Foxoa1, Hnf1a, Mapk15, Rgs16* and *Fasn.* RNA-seq data have the potential for application across diverse areas of human health conditions, including disease diagnosis, prognosis, and therapeutic selection (62). Understanding the mechanistic aspects of these genes, might give a clue for the early diagnosis and prognosis of hypertension.

This study has several limitations that should be acknowledged. hAGT cannot be readily cleaved by mouse renin. It requires co-expression of human renin to facilitate angiotensin II production. Crossing mice carrying human renin with -6A TG mice facilitates the production of angiotensinogen II. This would enable to study physiological functions such as blood pressure in -6A TG mice. However, in this study we focused on epigenetic aspect and transcriptional regulation of hAGT gene. In conclusion, while this study has contributed to our understanding of the epigenetic and transcriptional regulation of hAGT gene in the context of hypertension, it is important to acknowledge the limitations inherent in the study design.

## Conclusions and perspectives

To summarize, pathophysiological variables such as high salt modulates the DNA methylation pattern and transcriptional milieu that ultimately leads to the change in the hAGT gene expression. High-salt promotes DNA demethylation and increases the expression of inflammatory cytokines such as IL-6 and various transcription factors include C/EBPβ, GR, STAT-3, HNF-1α and USF1 in the liver and kidney tissues, which results in increased binding of these transcription factors to the open chromatin and increased expression of the hAGT gene and increased angiotensin II levels in -6A TG mice. This would potentially put people with -6A more susceptible to high salt-induced, AGT-mediated complications such as hypertension and other cardiovascular events. We have discovered the novel CpG sites on the hAGT promoter. And we also identified the CpG sites which are high-salt sensitive. Furthermore, we have identified several novel genes, pathways, upstream regulators implicated in the pathogenesis of hypertension. Clinical relevance of this study would be to help identify individuals with risk haplotypes. In this scenario, identification of “at-risk” haplotypes of the hAGT gene will assist patients by providing timely and targeted therapy to predict and prevent any long-term complications.

## Experimental procedures

### Generation of TG mice

The linearized plasmids were introduced into BK4 ES cells by electroporation and used to generate transgenic mice on the C57/BL6 background at Dartmouth Medical Center (Dartmouth, MA, USA) as described in previous literature (20, 63). Genotyping analysis of the tail snips, followed by sequencing, was performed to confirm the genetic lineage of these TG mice. These transgenic mice had a single copy of the hAGT gene, as determined by qPCR (64). All the animal experiments were performed according to the National Institutes of Health Guide for the Care and Use of Laboratory Animals and approved by the institutional ethical animal care and use committee at the University of Toledo College of Medicine and Life Sciences, Toledo, OH and New York Medical College, Valhalla, New York.

### High-salt (HS) diet

We gave 4% high salt to TG mice for 4 weeks to examine the effect of high salt on the DNA methylation pattern and the expression of the hAGT gene. Twelve weeks old adult male single transgenic mice containing -6A each were divided into two groups (n=6). Animals were maintained in a 22^0^ C room with a 12-hour light/dark cycle and received 4% HS and drinking water. Experiments were performed in the absence and presence of high salt.

### DNA methylation analysis

DNA methylation analysis was performed by EpiTect Bisulfite Kit (Cat.No.59104) from Qiagen. We isolated liver and kidney tissues from the -6A TG mice and isolated genomic DNA, converted gDNA to bisulfite DNA by using bisulfite kit by Qiagen. After bisulfite conversion all the methylated cytosines will be intact and un methylated cytosines are converted to (T) thymines. We amplified the bisulfite DNA from -483 to +105 region of the h AGT gene promoter by using -483 F and +105 R primers. Amplified DNA was run on agarose gel and purify DNA and clone the purified amplicon in TOPO TA vector. We prepared plasmid by using QIAprep Spin Miniprep Kit from Qiagen (Cat.No.27104 and Cat.No.27106). Plasmid was sequenced by using M13 R vector specific primers and analyzed the sequence and identified CpG sites by using bioinformatics software tools such as Vector NTI, Geneious Prime and BioEdit.

### RNA extraction and quantitative real-time PCR (qRT-PCR)

Liver and kidney tissues from 12 weeks old adult male transgenic mice were harvested at the end of the experiment following euthanasia. Snap frozen the liver and kidney tissues in liquid nitrogen immediately. Harvested tissues were further stored at -80 ^0^C until utilized for further experimental purpose. RNA was isolated using an RNeasy kit (Cat.No.74104) from QIAGEN. A total of 1 μg RNA was reverse transcribed to generate cDNA using a high-capacity cDNA reverse transcription kit (Cat.No.4368814) from Applied Biosystems, Thermo Fisher Scientific. qPCR was performed using PowerUp^TM^ SYBR^TM^ Green Master Mix (Cat.No. A25742; Applied Biosystems, Thermo Fisher Scientific) and the CFX Connect ^TM^ real-time PCR detection system (Bio-Rad). Primers for mouse AGT and human AGT were purchased from Integrated DNA Technologies and 18S rRNA primers were obtained from Keck Biotechnology Resource Laboratory. All relative gene expression levels were analyzed using the ΔΔC_T_ method and normalized to 18S rRNA expression.

### Immunoblot analysis

Blood samples were collected by cardiac puncture from mice immediately after exsanguination. Approximately 250μl plasma was collected from each mouse by centrifuging 500ul of blood sample at 3,000 rpm for 20 min at 4°C. Liver and kidney tissue lysates prepared from -6A basal and HS treated transgenic mice using RIPA lysis buffer. Liver and kidney tissue lysates were incubated at 4°C for 35 minutes and clarified by centrifugation at 14,000 rpm at 4°C for 30 minutes. Protein concentration was determined by using the Pierce^TM^ BCA Protein Assay Kit (Cat.No.23225; Thermo Fisher Scientific). Protein extracts (60 μg) were subjected to SDS-PAGE (10% polyacrylamide) and immunoblotting. After SDS-PAGE, gel transferred to 0.45 μm Immobilin-P (PVDF) transfer membranes to detect hAGT protein. The membranes were blocked for 1 hr with 1x TBS with 1% Casein (Cat.No.1610782; Bio-Rad) followed by overnight incubation at 4^0^ C with primary antibodies against human angiotensinogen (Cat.No.ab108294, 1:1000) from Abcam and albumin rabbit antibody (Cat.No.4929S; 1:1000) from cell signaling technology and beta-actin anti mouse mAb from Li-COR. We used rabbit and mouse secondary antibodies (1:30,000) conjugated with IRDye800 or IRDye600 for 60 minutes at room temperature protected from light with gentle shaking. Finally, all blots were visualized using the Odyssey CLx Imaging System (LI-COR Biosciences) and subjected to quantitative analyses using ImageJ software (version 1.5e). The results were normalized with albumin and beta-actin.

### In vivo chromatin immunoprecipitation (ChIP) assay

The chromatin immunoprecipitation (ChIP) assay was performed using the EZ-ChIP assay kit from EMD Millipore, MA, USA (56). Mice were perfused with normal saline. Liver and kidney tissues were excised and washed in 1X PBS, minced into smaller pieces; fixed with 1% formaldehyde for 25 mins at room temperature; washed with ice cold PBS followed by lysis. The DNA was fragmented by using sonication method and 10μl of the chromatin solution was saved as input. A 5μg of anti-glucocorticoid receptor (GR) or rabbit immunoglobulin G, anti Pol-II anti-STAT3, anti HNF1a, anti-USF1, anti C/EBP-#x2265; antibodies were added to the tubes containing 900μl of sonicated chromatin solution, and the mixture was incubated overnight at 4°C. The antibody complexes were captured with protein A-agarose beads and subjected to serial washes (as described in manufacturer’s protocol). The chromatin fraction was extracted with SDS buffer and reverse cross-linked at 65°C for 4-6 hrs.

The DNA was then purified as described in the manufacturer’s protocol. The immuno-precipitated DNA (1μl) and the input DNA (1μl) were subjected to PCR amplification (35 cycles: denaturation at 95^0^ C for 30 s, annealing at 58^0^ C for 30s, synthesis or extension at 72^0^ C for 30 s) using following primers (a) -217F AGT for (ATGCTCCCGTTTCTG GGAAC) as a forward and -217R AGT (CAGGCTGGAGAGGAGGGTTA) as a reverse primer and SYBR Green PCR Master Mix. Threshold cycles for three replicate reactions were determined, and relative enriched DNA abundance was calculated after normalization with input DNA.

### Transcriptome analysis in the liver and kidney of -6A TG mice

Total RNA was isolated from the liver and kidney of mice using an RNeasy kit (Cat.No.74104; QIAGEN) according to the manufacturer’s protocol. RNA quality was determined by estimating the A260/A280 and A260/A230 ratios by NanoDrop (Thermo Fisher Scientific). RNA integrity and quality was determined by running an Agilent Bioanalyzer gel, which measures the ratio of the ribosomal peaks. Statistical analyses were performed using Partek Flow software (version 7.0, build 7.18.0130.2018 Partek Inc., St Louis, Missouri). Paired-end reads were trimmed using a base quality score threshold of >24 and aligned to the Genome Reference Consortium Mouse Build 38 (mm10) with the STAR 2.5.3a aligner. Quantification was carried out using the transcript model Ensembl Transcripts release 100. Differential expression analysis was performed using the R package DESeq2 which provides a quantitative analysis using shrinkage estimators for dispersion and fold-change (Love et al., 2014). Genes in treated mice with an absolute fold-change of 1.5 and Benjamini-Hochberg false discovery rate (FDR) of p < 0.05 relative to control mice were considered as differentially expressed genes (DEGs). A negative fold-change indicates down-regulation of the treated mice compared with controls. Qlucore Omics Explorer 3.4 (Qlucore AB, Lund, Sweden) was used for data visualization, principal components analysis (PCA) and hierarchical clustering analysis (HCA) heatmap generation of global expression values. Gene Expression Omnibus database: GSE243600. Network analysis was performed using Ingenuity Pathway Analysis software (IPA, QIAGEN Redwood City, www.qiagen.com/ingenuity) to identify upstream regulators, pathways, diseases, and functions over-represented in the DEGs. In this manually curated knowledge base, each gene symbol was mapped to its corresponding gene object in the Ingenuity Pathway Knowledge Base. Significant (Benjamini-Hochberg FDR, p < 0.05) pathway enrichment within a reference network was performed using Fisher’s Exact test.

## Supporting information

Supporting Information_11_22_23 bioRxiv

## Statistical analysis

Statistical calculations with two-tailed Student’s t-test were done using GraphPad Prism software Version 10.0. All the data are presented are expressed as means with error bars representing standard error. p ≤ 0.05 was considered statistically significant.

## Data availability

The RNA-seq datasets mentioned in this article has been deposited in the NCBI Database of GEO Datasets under the accession number GSE243600.

## Supporting information

This article contains supporting information.

## Acknowledgments

We specially thank my mentor Ashok Kumar, Ph.D., FAHA, for his guidance and invaluable support. We thank Anton M. Bennett (Yale University), Meenakshi Kaw (University of Toledo), Sudhir Jain, Brahmaraju Mopidevi (New York Medical College), Shubham Mishra, Mansoor Ahmed, Jonathan Pascale (Yale University), Jyothi Podili (University of New Haven) for their technical support and critical discussions. We thank Yale Center for Genome Analysis (YCGA) for RNA Seq analysis. We specially thank Rolando Garcia Milan (Yale University) for his invaluable bioinformatics support. We also thank Nur-Taz Rahman (Yale University) for her suggestions regarding Qlucore Software.

## Author contributions

S.P. and A.K.: study concept and experimental design; S.P.: acquisition of data, analysis, interpretation of data, RNA-Seq analysis, manuscript preparation, preparation of figures and statistical analysis; A.K.: acquisition of funding and monitoring project; Both authors reviewed the manuscript.

## Funding and additional information

This work was supported by NHLBI, National Institutes of Health Grants RO1 HL 122742 and RO1 HL 130344 (to A.K.) Transgenic animals were generated at Dartmouth Medical Center. We acknowledge and appreciate help from the Dartmouth Mouse Modeling Shared Resource, which receives support from Norris Cotton Cancer Center Shared Grant P30CA023108.

