## Supporting Information_11_22_23 bioRxiv for "Epigenetic and transcriptional regulation of the human angiotensinogen gene by high salt": Supporting Information_11_22_23 bioRxiv.pdf

**Ashok Kumar, Ph.D., FAHA.**

Department of Pathology,

New York Medical College,

Valhalla, NY, USA, 10595.

.

**Table S1. Genes list from the liver by RNA Seq analysis.**

| High Salt vs Basal | | | |
|---|---|---|---|
| **LIVER** | | | |
| ***Gene symbol*** | **P-value** | **FDR step up** | **Fold Change** |
| *Fgf21* | 2.05E-06 | 0.000677 | 10.95 |
| *Nnmt* | 2.20E-07 | 2.17E-03 | 9.48 |
| *Hcn3* | 3.57E-07 | 2.17E-03 | 9.20 |
| *Rnf145* | 1.29E-06 | 2.96E-03 | 7.98 |
| *Rhbg* | 1.62E-06 | 2.96E-03 | 7.56 |
| *Cyp7b1* | 1.68E-06 | 2.96E-03 | 7.20 |
| *Rgs16* | 1.34E-08 | 1.02E-05 | 6.55 |
| *Acly* | 1.78E-06 | 2.96E-03 | 5.80 |
| *Capn8* | 1.95E-06 | 2.96E-03 | 5.34 |
| *Pklr* | 4.26E-06 | 5.44E-03 | 4.83 |
| *Selenbp2* | 5.33E-06 | 5.44E-03 | 4.73 |
| *Fasn* | 5.36E-06 | 5.44E-03 | 4.68 |
| *Elovl5* | 5.74E-06 | 5.44E-03 | 4.57 |
| *Lpar2* | 5.81E-06 | 5.44E-03 | 4.03 |
| *Acaca* | 8.08E-06 | 7.02E-03 | 3.90 |
| *Rpia* | 1.18E-05 | 9.33E-03 | 3.90 |
| *Scd1* | 1.23E-05 | 9.33E-03 | 3.81 |
| *Ltbp4* | 1.38E-05 | 9.90E-03 | 3.81 |
| *Dio1* | 1.57E-05 | 1.04E-02 | 3.77 |
| *Dsp* | 1.62E-05 | 1.04E-02 | 3.76 |
| *Tmem131l* | 1.73E-05 | 1.04E-02 | 3.62 |
| *Eef2k* | 1.97E-05 | 1.04E-02 | 3.47 |
| *Me1* | 2.06E-05 | 1.04E-02 | 3.45 |
| *Mapk15* | 3.43E-07 | 0.000162 | 3.36 |
| *Mup18* | 2.14E-05 | 1.04E-02 | 3.31 |
| *Dll1* | 2.15E-05 | 1.04E-02 | 3.30 |
| *Cyp46a1* | 2.23E-05 | 1.04E-02 | 3.26 |
| *Elovl6* | 2.39E-05 | 1.07E-02 | 3.23 |
| *Sult5a1* | 2.52E-05 | 1.09E-02 | 3.21 |
| *Fnip2* | 2.75E-05 | 1.12E-02 | 3.13 |
| *Pltp* | 2.77E-05 | 1.12E-02 | 3.09 |
| *Mup17* | 3.03E-05 | 1.13E-02 | 3.01 |
| *Iffo2* | 3.16E-05 | 1.13E-02 | 2.96 |
| *Als2* | 3.36E-05 | 1.17E-02 | 2.77 |
| *Derl3* | 5.98E-05 | 2.02E-02 | 2.76 |
| *Syne3* | 6.73E-05 | 2.21E-02 | 2.73 |
| *Chid1* | 7.80E-05 | 2.50E-02 | 2.68 |
| *Svil* | 8.32E-05 | 2.60E-02 | 2.67 |
| *Hnf1a* | 1.12E-06 | 0.000419 | 2.66 |
| *Chrna4* | 9.36E-05 | 2.72E-02 | 2.61 |
| *Hnrnpd* | 9.38E-05 | 2.72E-02 | 2.58 |
| *Tmem62* | 1.02E-04 | 2.78E-02 | 2.58 |

| | | | |
|---|---|---|---|
| *Tmie* | 1.05E-04 | 2.78E-02 | 2.55 |
| *Dapk2* | 1.05E-04 | 2.78E-02 | 2.55 |
| *Arsa* | 1.07E-04 | 2.78E-02 | 2.54 |
| *Ttl* | 1.07E-04 | 2.78E-02 | 2.54 |
| *Obp2a* | 1.17E-04 | 2.96E-02 | 2.48 |
| *Arl4a* | 1.22E-04 | 3.02E-02 | 2.43 |
| *Cyp2b10* | 1.26E-04 | 3.03E-02 | 2.39 |
| *Reck* | 1.29E-04 | 3.03E-02 | 2.38 |
| *Spata13* | 1.30E-04 | 3.03E-02 | 2.33 |
| *Slc35b1* | 1.32E-04 | 3.03E-02 | 2.32 |
| *Acmsd* | 1.36E-04 | 3.06E-02 | 2.29 |
| *Rassf6* | 1.43E-04 | 3.16E-02 | 2.24 |
| *Sdf2l1* | 1.46E-04 | 3.17E-02 | 2.22 |
| *Ugt1a5* | 1.53E-04 | 3.27E-02 | 2.20 |
| *Tkfc* | 1.65E-04 | 3.40E-02 | 2.20 |
| *C9* | 1.68E-04 | 3.41E-02 | 2.12 |
| *Gsn* | 1.77E-04 | 3.49E-02 | 2.12 |
| *Pgap2* | 1.78E-04 | 3.49E-02 | 2.12 |
| *Pxmp2* | 1.83E-04 | 3.52E-02 | 2.11 |
| *Rnf152* | 1.85E-04 | 3.52E-02 | 2.07 |
| *Bahcc1* | 1.91E-04 | 3.57E-02 | 2.06 |
| *Steap2* | 2.09E-04 | 3.66E-02 | 2.04 |
| *Asic5* | 2.10E-04 | 3.66E-02 | 2.02 |
| *Gpi1* | 2.13E-04 | 3.66E-02 | 1.98 |
| *Snx22* | 2.13E-04 | 3.66E-02 | 1.97 |
| *Vwce* | 2.14E-04 | 3.66E-02 | 1.90 |
| *Smim1* | 2.21E-04 | 3.66E-02 | 1.89 |
| *Mup8* | 2.23E-04 | 3.66E-02 | 1.88 |
| *Als2cl* | 2.27E-04 | 3.66E-02 | 1.86 |
| *Hlf* | 2.31E-04 | 3.66E-02 | 1.85 |
| *Aig1* | 2.33E-04 | 3.66E-02 | 1.84 |
| *Gna14* | 2.37E-04 | 3.66E-02 | 1.83 |
| *Zfp385a* | 2.40E-04 | 3.66E-02 | 1.81 |
| *Lysmd4* | 2.40E-04 | 3.66E-02 | 1.81 |
| *Hipk2* | 2.41E-04 | 3.66E-02 | 1.74 |
| *Tns3* | 2.50E-04 | 3.74E-02 | 1.73 |
| *Nucb2* | 2.53E-04 | 3.74E-02 | 1.72 |
| *Hspb8* | 2.55E-04 | 3.74E-02 | 1.68 |
| *Per3* | 2.60E-04 | 3.74E-02 | 1.64 |
| *Morn2* | 2.71E-04 | 3.83E-02 | -1.65 |
| *Tmem120a* | 2.81E-04 | 3.89E-02 | -1.74 |
| *Smurf1* | 2.81E-04 | 3.89E-02 | -1.74 |
| *Frmd6* | 2.84E-04 | 3.89E-02 | -1.74 |
| *Ddhd2* | 2.89E-04 | 3.91E-02 | -1.80 |
| *Lzts2* | 2.95E-04 | 3.93E-02 | -1.82 |
| *Tfcp2l1* | 2.97E-04 | 3.93E-02 | -1.82 |
| *Pccb* | 3.10E-04 | 3.98E-02 | -1.84 |

| | | | |
|---|---|---|---|
| *Mup13* | 3.12E-04 | 3.98E-02 | -1.85 |
| *Enpep* | 3.15E-04 | 3.98E-02 | -1.88 |
| *Zbtb12* | 3.18E-04 | 3.98E-02 | -1.88 |
| *Fmo4* | 3.32E-04 | 4.12E-02 | -1.91 |
| *Pou2f1* | 3.41E-04 | 4.16E-02 | -1.92 |
| *Msmo1* | 3.42E-04 | 4.16E-02 | -1.96 |
| *Gstm2* | 3.65E-04 | 4.21E-02 | -2.01 |
| *Plbd1* | 3.69E-04 | 4.21E-02 | -2.07 |
| *Cyp3a13* | 3.71E-04 | 4.21E-02 | -2.13 |
| *Unc5b* | 3.73E-04 | 4.21E-02 | -2.14 |
| *Vegfb* | 3.75E-04 | 4.21E-02 | -2.18 |
| *Lrrc28* | 3.79E-04 | 4.21E-02 | -2.18 |
| *Inhba* | 3.80E-04 | 4.21E-02 | -2.21 |
| *Acacb* | 3.84E-04 | 4.21E-02 | -2.22 |
| *Enho* | 3.87E-04 | 4.21E-02 | -2.36 |
| *Aqp8* | 3.96E-04 | 4.26E-02 | -2.40 |
| *Gla* | 4.08E-04 | 4.30E-02 | -2.41 |
| *Actn1* | 4.09E-04 | 4.30E-02 | -2.41 |
| *Bcl2l11* | 4.14E-04 | 4.30E-02 | -2.45 |
| *Rcl1* | 4.14E-04 | 4.30E-02 | -2.45 |
| *Agxt2* | 4.18E-04 | 4.31E-02 | -2.53 |
| *Sox12* | 4.29E-04 | 4.38E-02 | -2.60 |
| *Irs1* | 4.47E-04 | 4.46E-02 | -2.68 |
| *Mup14* | 4.51E-04 | 4.46E-02 | -2.72 |
| *Crat* | 4.51E-04 | 4.46E-02 | -2.92 |
| *Cyp1a2* | 4.55E-04 | 4.47E-02 | -2.96 |
| *Rnd1* | 4.63E-04 | 4.51E-02 | -2.96 |
| *Kank2* | 4.68E-04 | 4.52E-02 | -2.97 |
| *Oat* | 4.73E-04 | 4.53E-02 | -3.03 |
| *Eef1akmt4* | 4.81E-04 | 4.54E-02 | -3.06 |
| *F11* | 4.89E-04 | 4.54E-02 | -3.55 |
| *Dcun1d5* | 4.90E-04 | 4.54E-02 | -3.56 |
| *Eea1* | 4.90E-04 | 4.54E-02 | -3.58 |
| *Sparcl1* | 4.92E-04 | 4.54E-02 | -3.69 |
| *Wfdc21* | 5.12E-04 | 4.65E-02 | -3.85 |
| *Nfic* | 5.12E-04 | 4.65E-02 | -4.07 |
| *Inca1* | 5.17E-04 | 4.65E-02 | -4.20 |
| *Phf2* | 5.20E-04 | 4.65E-02 | -4.48 |
| *Tango2* | 5.37E-04 | 4.70E-02 | -4.92 |
| *Wwc2* | 5.38E-04 | 4.70E-02 | -5.01 |
| *Bhmt* | 5.41E-04 | 4.70E-02 | -5.27 |
| *Hnf1b* | 5.42E-04 | 4.70E-02 | -5.90 |
| *Itpa* | 5.44E-04 | 4.70E-02 | -7.21 |

**Table S2. Genes list from the kidney by RNA Seq analysis.**

| High Salt vs Basal | | |
|---|---|---|
| **KIDNEY** | | |
| ***Gene symbol*** | **P-value** | **Fold Change** |
| *Nnmt* | 2.20E-07 | 9.48 |
| *Gm4322* | 1.40E-07 | 14.11 |
| *Gm13855* | 4.92E-07 | 13.67 |
| *Myh6* | 5.42E-07 | 9.22 |
| *Hrg* | 6.16E-07 | 6.91 |
| *Xlr4b* | 5.93E-06 | 6.44 |
| *Serpina3k* | 8.93E-06 | 6.39 |
| *Ttn* | 1.21E-05 | 5.87 |
| *Gm48899* | 2.06E-05 | 5.33 |
| *Gm1966* | 2.24E-05 | 5.30 |
| *Srd5a1* | 2.35E-05 | 5.28 |
| *Cyp2a5* | 2.41E-05 | 4.98 |
| *Abcb1b* | 5.45E-05 | 4.97 |
| *Dock4* | 5.91E-05 | 4.56 |
| *Gm13841* | 1.07E-04 | 4.48 |
| *S100a9* | 1.33E-04 | 4.38 |
| *Gm15590* | 1.94E-04 | 4.33 |
| *Gm13233* | 2.26E-04 | 4.32 |
| *Gm9108* | 2.52E-04 | 4.25 |
| *Htatip2* | 3.29E-04 | 4.15 |
| *Gm9493* | 4.19E-04 | 4.09 |
| *Aldh1a7* | 4.46E-04 | 4.04 |
| *A230056P14Rik* | 5.63E-04 | 3.99 |
| *Gm20427* | 6.83E-04 | 3.88 |
| *Apoh* | 8.34E-04 | 3.87 |
| *Mogat2* | 9.43E-04 | 3.86 |
| *Ctse* | 9.43E-04 | 3.85 |
| *Hapln3* | 1.08E-03 | 3.79 |
| *Ptger3* | 1.17E-03 | 3.77 |
| *Tpi-rs10* | 1.19E-03 | 3.66 |
| *Gm42047* | 1.19E-03 | 3.61 |
| *Prlr* | 1.32E-03 | 3.44 |
| *Fcgbp* | 1.40E-03 | 3.40 |
| *Gm20708* | 1.41E-03 | 3.32 |
| *Gm6109* | 1.41E-03 | 3.29 |
| *Gm5276* | 1.55E-03 | 3.27 |
| *Gfra1* | 1.64E-03 | 3.25 |
| *Wfdc16* | 1.78E-03 | 3.19 |
| *Arntl2* | 1.82E-03 | 3.18 |
| *Cadm2* | 1.83E-03 | 3.17 |
| *Ren1* | 1.89E-03 | 3.14 |
| *Gm17802* | 1.97E-03 | 3.12 |
| *Gm3776* | 2.14E-03 | 3.10 |

| | | |
|---|---|---|
| *Rps4l* | 2.27E-03 | 3.08 |
| *Atp4a* | 2.37E-03 | 3.06 |
| *Aldoart2* | 2.40E-03 | 2.98 |
| *Cd209a* | 2.57E-03 | 2.96 |
| *Ctxn3* | 2.60E-03 | 2.94 |
| *Hmgb1-ps3* | 3.07E-03 | 2.94 |
| *Gm6663* | 3.08E-03 | 2.92 |
| *Tm4sf4* | 3.08E-03 | 2.88 |
| *Hspa1b* | 3.14E-03 | 2.88 |
| *Brms1* | 3.32E-03 | 2.86 |
| *Aldh1a1* | 3.38E-03 | 2.85 |
| *Chn2* | 3.43E-03 | 2.84 |
| *Sptssb* | 3.89E-03 | 2.74 |
| *Klk1b16* | 3.98E-03 | 2.74 |
| *Kcnma1* | 4.13E-03 | 2.73 |
| *Ighm* | 4.14E-03 | 2.69 |
| *Spcs2-ps* | 4.15E-03 | 2.69 |
| *Mup20* | 4.19E-03 | 2.65 |
| *Ftl1* | 4.31E-03 | 2.62 |
| *1810064F22Rik* | 4.41E-03 | 2.62 |
| *Gm6394* | 4.59E-03 | 2.61 |
| *Tmem178b* | 5.03E-03 | 2.61 |
| *Gpr183* | 5.10E-03 | 2.58 |
| *C130012C08Rik* | 5.14E-03 | 2.55 |
| *Gm47248* | 5.16E-03 | 2.51 |
| *Gm21092* | 5.29E-03 | 2.51 |
| *Gm37988* | 5.40E-03 | 2.49 |
| *Gm7935* | 5.41E-03 | 2.49 |
| *Gm50388* | 5.42E-03 | 2.47 |
| *Gm11400* | 5.48E-03 | 2.45 |
| *Hspa9-ps1* | 5.65E-03 | 2.45 |
| *Ldha-ps2* | 5.67E-03 | 2.44 |
| *Fam43a* | 5.70E-03 | 2.43 |
| *Gm8130* | 5.83E-03 | 2.42 |
| *Gm4617* | 5.88E-03 | 2.41 |
| *Gm4707* | 6.16E-03 | 2.40 |
| *Syk* | 6.25E-03 | 2.38 |
| *Cyp3a11* | 6.26E-03 | 2.38 |
| *Gm906* | 6.27E-03 | 2.37 |
| *Hspa1a* | 6.28E-03 | 2.35 |
| *Gm4471* | 6.30E-03 | 2.35 |
| *Igsf9b* | 6.35E-03 | 2.32 |
| *6330410L21Rik* | 6.72E-03 | 2.32 |
| *Fam177a* | 6.80E-03 | 2.29 |
| *Tmem151b* | 6.83E-03 | 2.29 |
| *Capg* | 6.84E-03 | 2.27 |
| *Lox* | 6.86E-03 | 2.27 |

| | | |
|---|---|---|
| *4930405A21Rik* | 7.15E-03 | 2.26 |
| *Gm15502* | 7.43E-03 | 2.24 |
| *Gm13772* | 7.57E-03 | 2.21 |
| *Trim12a* | 7.59E-03 | 2.21 |
| *Gvin1* | 7.70E-03 | 2.20 |
| *Dlg2* | 7.75E-03 | 2.20 |
| *Hddc3* | 8.02E-03 | 2.19 |
| *Chsy1* | 8.20E-03 | 2.18 |
| *Gm5881* | 8.62E-03 | 2.18 |
| *Clca3a1* | 8.84E-03 | 2.18 |
| *Anks1b* | 8.86E-03 | 2.17 |
| *A2m* | 8.89E-03 | 2.16 |
| *Cyp2b10* | 9.00E-03 | 2.15 |
| *Gm7392* | 9.01E-03 | 2.15 |
| *Acnat2* | 9.19E-03 | 2.14 |
| *Trim34b* | 9.21E-03 | 2.14 |
| *Gm9732* | 9.23E-03 | 2.14 |
| *Slc38a4* | 9.54E-03 | 2.12 |
| *Zfp935* | 9.65E-03 | 2.11 |
| *Acy1* | 9.82E-03 | 2.09 |
| *Tnnt2* | 9.90E-03 | 2.08 |
| *Snhg14* | 9.96E-03 | 2.07 |
| *Trdn* | 1.00E-02 | 2.06 |
| *Gm27029* | 1.00E-02 | 2.06 |
| *Gm8834* | 1.01E-02 | 2.05 |
| *Gm9315* | 1.05E-02 | 2.05 |
| *Gm10259* | 1.07E-02 | 2.04 |
| *Ccng1* | 1.13E-02 | 2.03 |
| *Gm14121* | 1.14E-02 | 2.03 |
| *D330041H03Rik* | 1.14E-02 | 2.02 |
| *Gm34220* | 1.14E-02 | 2.02 |
| *Nepn* | 1.14E-02 | 2.00 |
| *Gm48362* | 1.16E-02 | 2.00 |
| *Cyp24a1* | 1.19E-02 | 1.98 |
| *Gm8099* | 1.20E-02 | 1.98 |
| *Pvalb* | 1.21E-02 | 1.97 |
| *Parp6* | 1.21E-02 | 1.97 |
| *Gm7407* | 1.24E-02 | 1.96 |
| *Rpl28-ps3* | 1.25E-02 | 1.96 |
| *Rps7-ps2* | 1.25E-02 | 1.95 |
| *Gm9780* | 1.29E-02 | 1.95 |
| *Zfp457* | 1.29E-02 | 1.94 |
| *Ttc38* | 1.31E-02 | 1.93 |
| *Gm11964* | 1.32E-02 | 1.93 |
| *Gm16470* | 1.33E-02 | 1.92 |
| *Eda2r* | 1.35E-02 | 1.91 |
| *Fgb* | 1.36E-02 | 1.91 |

| | | |
|---|---|---|
| *Gm47283* | 1.39E-02 | 1.90 |
| *Ldha* | 1.39E-02 | 1.89 |
| *Gm5586* | 1.41E-02 | 1.89 |
| *Arg2* | 1.41E-02 | 1.89 |
| *Rps3a3* | 1.43E-02 | 1.88 |
| *Cndp1* | 1.43E-02 | 1.88 |
| *Cdc42ep2* | 1.43E-02 | 1.87 |
| *Gjb1* | 1.45E-02 | 1.86 |
| *Slc28a1* | 1.48E-02 | 1.85 |
| *Lypd6* | 1.49E-02 | 1.85 |
| *Irf3* | 1.51E-02 | 1.83 |
| *Gm10399* | 1.55E-02 | 1.83 |
| *Kcnj2* | 1.56E-02 | 1.82 |
| *Kcnk2* | 1.56E-02 | 1.82 |
| *Gm4973* | 1.58E-02 | 1.81 |
| *Plvap* | 1.59E-02 | 1.80 |
| *Gabrb3* | 1.59E-02 | 1.79 |
| *Ifi44l* | 1.62E-02 | 1.78 |
| *Gm49320* | 1.63E-02 | 1.78 |
| *Angpt2* | 1.66E-02 | 1.78 |
| *Pear1* | 1.67E-02 | 1.78 |
| *Ttyh1* | 1.68E-02 | 1.76 |
| *Gm12096* | 1.70E-02 | 1.76 |
| *B230303O12Rik* | 1.72E-02 | 1.76 |
| *Unc5c* | 1.73E-02 | 1.76 |
| *Gm6532* | 1.73E-02 | 1.76 |
| *Gm16378* | 1.75E-02 | 1.75 |
| *Gm37245* | 1.75E-02 | 1.75 |
| *Uox* | 1.76E-02 | 1.74 |
| *Oasl2* | 1.76E-02 | 1.73 |
| *Fabp1* | 1.77E-02 | 1.73 |
| *Gm8319* | 1.80E-02 | 1.73 |
| *Gm38317* | 1.82E-02 | 1.72 |
| *Lrrc17* | 1.83E-02 | 1.72 |
| *Acat3* | 1.84E-02 | 1.71 |
| *Gm6813* | 1.84E-02 | 1.70 |
| *Gm10130* | 1.85E-02 | 1.68 |
| *Cep85* | 1.85E-02 | 1.67 |
| *Gm10108* | 1.86E-02 | 1.67 |
| *Tenm4* | 1.86E-02 | 1.66 |
| *Il12b* | 1.87E-02 | 1.65 |
| *Gm47428* | 1.88E-02 | 1.65 |
| *Gm8349* | 1.90E-02 | 1.64 |
| *Gm31651* | 1.91E-02 | 1.63 |
| *Gm11196* | 1.93E-02 | 1.62 |
| *Upk3b* | 1.95E-02 | 1.62 |
| *Syt17* | 1.99E-02 | 1.61 |

| | | |
|---|---|---|
| *Gm7329* | 2.02E-02 | 1.60 |
| *Cxcr3* | 2.02E-02 | 1.60 |
| *Morn4* | 2.04E-02 | 1.60 |
| *Ciart* | 2.06E-02 | 1.59 |
| *C4bp* | 2.09E-02 | 1.59 |
| *Procr* | 2.11E-02 | 1.58 |
| *Hapln1* | 2.11E-02 | 1.57 |
| *AW822252* | 2.14E-02 | 1.56 |
| *Tubgcp5* | 2.16E-02 | 1.55 |
| *Rpl21-ps11* | 2.16E-02 | 1.55 |
| *Gm36210* | 2.17E-02 | 1.54 |
| *BC085271* | 2.17E-02 | 1.53 |
| *Zfp966* | 2.19E-02 | 1.53 |
| *8430426J06Rik* | 2.21E-02 | 1.50 |
| *Arhgap25* | 2.24E-02 | -1.50 |
| *Gstp2* | 2.26E-02 | -1.52 |
| *Gm29737* | 2.27E-02 | -1.52 |
| *Gm8738* | 2.29E-02 | -1.53 |
| *Loxl4* | 2.30E-02 | -1.54 |
| *Slc23a3* | 2.30E-02 | -1.58 |
| *Gm5513* | 2.31E-02 | -1.58 |
| *Trim34a* | 2.32E-02 | -1.61 |
| *Rgs4* | 2.34E-02 | -1.62 |
| *A930038B10Rik* | 2.37E-02 | -1.63 |
| *Hhatl* | 2.37E-02 | -1.65 |
| *C230037L18Rik* | 2.41E-02 | -1.66 |
| *Clec10a* | 2.47E-02 | -1.66 |
| *Ppp1r2-ps1* | 2.51E-02 | -1.67 |
| *Ly6d* | 2.55E-02 | -1.67 |
| *Znrd2* | 2.57E-02 | -1.67 |
| *Prickle2* | 2.58E-02 | -1.68 |
| *Lrat* | 2.58E-02 | -1.68 |
| *H3f3aos* | 2.58E-02 | -1.68 |
| *Cd79a* | 2.60E-02 | -1.68 |
| *Zfp61* | 2.61E-02 | -1.68 |
| *4932422M17Rik* | 2.66E-02 | -1.69 |
| *Pcdhb11* | 2.71E-02 | -1.69 |
| *Gm45053* | 2.74E-02 | -1.70 |
| *Gli1* | 2.76E-02 | -1.71 |
| *Bcl2a1a* | 2.78E-02 | -1.71 |
| *Abcg3* | 2.78E-02 | -1.73 |
| *Gm5518* | 2.78E-02 | -1.75 |
| *Gm11225* | 2.81E-02 | -1.75 |
| *Gm8483* | 2.83E-02 | -1.76 |
| *Phtf1os* | 2.85E-02 | -1.77 |
| *Gm38393* | 2.85E-02 | -1.77 |
| *Gm8784* | 2.85E-02 | -1.81 |

| | | |
|---|---|---|
| *Kcnrg* | 2.88E-02 | -1.82 |
| *B3gnt2* | 2.88E-02 | -1.82 |
| *Il1b* | 2.91E-02 | -1.83 |
| *Gm16685* | 2.92E-02 | -1.83 |
| *Ptger4* | 2.94E-02 | -1.83 |
| *Gm10138* | 2.95E-02 | -1.84 |
| *Gm15567* | 2.95E-02 | -1.85 |
| *Cyp4a10* | 2.96E-02 | -1.85 |
| *Npl* | 2.99E-02 | -1.85 |
| *A830082K12Rik* | 3.00E-02 | -1.86 |
| *Ces1c* | 3.00E-02 | -1.86 |
| *Zfp791* | 3.01E-02 | -1.88 |
| *Gm11625* | 3.03E-02 | -1.90 |
| *Coch* | 3.08E-02 | -1.90 |
| *B230377A18Rik* | 3.11E-02 | -1.90 |
| *Cyp4a14* | 3.12E-02 | -1.92 |
| *Gm5131* | 3.12E-02 | -1.93 |
| *Ggct* | 3.12E-02 | -1.93 |
| *Lax1* | 3.12E-02 | -1.94 |
| *Ccne2* | 3.13E-02 | -1.97 |
| *Gm4724* | 3.13E-02 | -2.00 |
| *Rps13-ps1* | 3.16E-02 | -2.00 |
| *Fjx1* | 3.16E-02 | -2.01 |
| *Gm12612* | 3.16E-02 | -2.04 |
| *Rsad1* | 3.19E-02 | -2.04 |
| *Egln3* | 3.22E-02 | -2.04 |
| *Gm49594* | 3.25E-02 | -2.05 |
| *Gm9029* | 3.26E-02 | -2.05 |
| *Gm17819* | 3.26E-02 | -2.05 |
| *Eif5al3-ps* | 3.27E-02 | -2.06 |
| *Cyp3a13* | 3.36E-02 | -2.11 |
| *Gm12940* | 3.36E-02 | -2.12 |
| *Atp5g2* | 3.37E-02 | -2.13 |
| *Col13a1* | 3.39E-02 | -2.17 |
| *Tmem207* | 3.41E-02 | -2.17 |
| *A730020M07Rik* | 3.44E-02 | -2.17 |
| *Id1* | 3.47E-02 | -2.19 |
| *2810454H06Rik* | 3.47E-02 | -2.23 |
| *Blvrb* | 3.48E-02 | -2.29 |
| *Abhd8* | 3.49E-02 | -2.31 |
| *Zfp992* | 3.50E-02 | -2.31 |
| *Ccl21b* | 3.51E-02 | -2.35 |
| *Gm38118* | 3.54E-02 | -2.37 |
| *Enox1* | 3.56E-02 | -2.40 |
| *Aplnr* | 3.56E-02 | -2.42 |
| *Spats1* | 3.60E-02 | -2.42 |
| *Gm2022* | 3.64E-02 | -2.42 |

| | | |
|---|---|---|
| *Wfdc17* | 3.65E-02 | -2.45 |
| *Gm50387* | 3.66E-02 | -2.46 |
| *Chst3* | 3.66E-02 | -2.47 |
| *BC064078* | 3.67E-02 | -2.49 |
| *Gm6135* | 3.67E-02 | -2.50 |
| *Tlr12* | 3.69E-02 | -2.51 |
| *Gm46379* | 3.70E-02 | -2.52 |
| *Gm2115* | 3.72E-02 | -2.53 |
| *Adgrg5* | 3.73E-02 | -2.53 |
| *U2af1* | 3.77E-02 | -2.53 |
| *Gm20369* | 3.77E-02 | -2.53 |
| *Irx5* | 3.82E-02 | -2.54 |
| *Gm37829* | 3.82E-02 | -2.56 |
| *Syt9* | 3.84E-02 | -2.57 |
| *Mup3* | 3.88E-02 | -2.57 |
| *Gm16170* | 3.90E-02 | -2.59 |
| *Hrct1* | 3.91E-02 | -2.62 |
| *Gm14928* | 3.91E-02 | -2.63 |
| *Gm17828* | 3.92E-02 | -2.63 |
| *Gm12732* | 3.93E-02 | -2.64 |
| *Tigd5* | 3.95E-02 | -2.66 |
| *Srcin1* | 3.98E-02 | -2.67 |
| *Gm49314* | 4.02E-02 | -2.68 |
| *2010003K11Rik* | 4.02E-02 | -2.72 |
| *A730049H05Rik* | 4.02E-02 | -2.73 |
| *Tdg-ps2* | 4.03E-02 | -2.74 |
| *mt-Rnr2* | 4.04E-02 | -2.74 |
| *Tmc8* | 4.05E-02 | -2.76 |
| *1110019B22Rik* | 4.05E-02 | -2.78 |
| *Atad3aos* | 4.05E-02 | -2.80 |
| *Gm42397* | 4.08E-02 | -2.81 |
| *Gm9619* | 4.09E-02 | -2.85 |
| *mt-Co1* | 4.10E-02 | -2.92 |
| *Gm14226* | 4.10E-02 | -2.94 |
| *Ccdc150* | 4.12E-02 | -3.08 |
| *Gm14328* | 4.12E-02 | -3.08 |
| *Gm6344* | 4.14E-02 | -3.08 |
| *Hpd* | 4.15E-02 | -3.18 |
| *Gm7977* | 4.16E-02 | -3.25 |
| *Gm10509* | 4.16E-02 | -3.27 |
| *Cenpk* | 4.18E-02 | -3.32 |
| *Gm5256* | 4.18E-02 | -3.37 |
| *Hmcn2* | 4.19E-02 | -3.44 |
| *Gm31300* | 4.24E-02 | -3.44 |
| *Vmn1r184* | 4.30E-02 | -3.49 |
| *Gm4890* | 4.31E-02 | -3.57 |
| *Gpr85* | 4.31E-02 | -3.59 |

| | | |
|---|---|---|
| *Gm13578* | 4.34E-02 | -3.60 |
| *Eif2s3x-ps1* | 4.35E-02 | -3.60 |
| *Gm2016* | 4.35E-02 | -3.60 |
| *6430511E19Rik* | 4.35E-02 | -3.72 |
| *Krt8* | 4.39E-02 | -3.72 |
| *Fam189a2* | 4.39E-02 | -3.78 |
| *AC154200.1* | 4.41E-02 | -3.78 |
| *Gm13196* | 4.44E-02 | -3.82 |
| *Hyal3* | 4.47E-02 | -3.82 |
| *Mir6236* | 4.49E-02 | -3.88 |
| *Cideb* | 4.50E-02 | -3.94 |
| *Gm30162* | 4.50E-02 | -3.94 |
| *C730036E19Rik* | 4.51E-02 | -3.95 |
| *Hspb1* | 4.52E-02 | -4.13 |
| *Rpl15-ps6* | 4.52E-02 | -4.25 |
| *Cct3-ps1* | 4.52E-02 | -4.25 |
| *B230322F03Rik* | 4.53E-02 | -4.26 |
| *Xlr3b* | 4.57E-02 | -4.29 |
| *Gm20045* | 4.59E-02 | -4.38 |
| *Gm34084* | 4.62E-02 | -4.53 |
| *Gm16373* | 4.63E-02 | -4.56 |
| *Alb* | 4.64E-02 | -4.64 |
| *Il2rg* | 4.66E-02 | -5.03 |
| *Gm10075* | 4.67E-02 | -5.05 |
| *Gm22513* | 4.69E-02 | -5.24 |
| *Dpp10* | 4.72E-02 | -5.29 |
| *Gm7560* | 4.73E-02 | -5.53 |
| *Cdr2l* | 4.74E-02 | -5.54 |
| *Nqo1* | 4.75E-02 | -5.56 |
| *Rpl29* | 4.75E-02 | -5.62 |
| *Mt2* | 4.76E-02 | -5.83 |
| *Lamc3* | 4.77E-02 | -6.08 |
| *Dnm1* | 4.78E-02 | -6.30 |
| *Gm45792* | 4.79E-02 | -6.76 |
| *Gm15411* | 4.81E-02 | -7.06 |
| *Pkdrej* | 4.85E-02 | -7.34 |
| *Aqp4* | 4.85E-02 | -7.42 |
| *Vars* | 4.85E-02 | -7.49 |
| *Mrto4-ps2* | 4.86E-02 | -8.09 |
| *Gm18705* | 4.86E-02 | -9.20 |
| *C030037D09Rik* | 4.89E-02 | -9.57 |
| *Gm17753* | 4.89E-02 | -9.59 |
| *Gm36936* | 4.92E-02 | -11.83 |
| *Gm37335* | 4.93E-02 | -12.67 |
| *Gm5778* | 4.95E-02 | -12.81 |
| *Mtbp* | 4.95E-02 | -15.55 |
| *Smim10l2a* | 4.96E-02 | -17.29 |

| | | |
|---|---|---|
| *Itga7* | 4.98E-02 | -31.09 |

**Table S3. The list of primers for quantitative real-time PCR analysis.**

| Primer Name | Sequences |
|---|---|
| *18S rRNA* | 5'-ACCGCAGCTAGGAATAATGGA-3'<br>5'-ACCAAAAGCCTTGACTCCG-3' |
| *mGAPDH* | 5'-CATCACTGCCACCCAGAAGACTG-5'<br>5'-ATGCCAGTGAGCTTCCCGTTCAG-3' |
| *hAGT* | 5'-TGGACAGCACCCTGGCTTTCAA-3'<br>5'-ACACTGAGGTGCTGTTGTCCAC-3' |
| *mAGT* | 5'-GGTCAGTACAGACAGCACCCTA-3'<br>5'-ACACCGAGATGCTGTTGTCCAC-3' |
| *Fasn* | *5'-CACAGTGCTCAAAGGACATGCC-3'*<br>*5'-CACCAGGTGTAGTGCCTTCCTC-3'* |
| *Fgf21* | *5'-ATCAGGGAGGATGGAACAGTGG-3'*<br>*5'-AGCTCCATCTGGCTGTTGGCAA-3'* |
| *Mapk15* | *5'-TCCAGGACCTTGGCTCAGACTA-3'*<br>*5'-AGCAAACGCCAAGAGTCGCTTG-3'* |
| *Rgs16* | *5'-ATCCGATCAGCCACCAAACTGG-3'*<br>*5'-GAAGCAACTGGTAGTGGCAGCT-3'* |
| *Hnf1a* | *5'-AGAGACCTTGGTGGAGGAGTGT-3'*<br>*5'-GGCAAACCAGTTGTAGACACGC-3'* |
